# Phylogenies of extant species are consistent with an infinite array of diversification histories

**DOI:** 10.1101/719435

**Authors:** Stilianos Louca, Matthew W. Pennell

**Author notes:** Correspondence should be addressed to both authors: www.loucalab.com and.

## Abstract

Time-calibrated molecular phylogenies of extant species ("extant timetrees") are widely used for estimating the dynamics of diversification rates (1–6) and testing for associations between these rates and environmental factors (5, 7) or species traits (8). However, there has been considerable debate surrounding the reliability of these inferences in the absence of fossil data (9–13), and to date this critical question remains unresolved. Here we mathematically clarify the precise information that can be extracted from extant timetrees under the generalized birth-death model, which underlies the majority of existing estimation methods. We prove that for a given extant timetree and a candidate diversification scenario, there exists an infinite number of alternative diversification scenarios that are equally likely to have generated a given tree. These “congruent” scenarios cannot possibly be distinguished using extant timetrees alone, even in the presence of infinite data. Importantly, congruent diversification scenarios can exhibit markedly different and yet plausible diversification dynamics, suggesting that many previous studies may have over-interpreted phylogenetic evidence. We show that sets of congruent models can be uniquely described using composite variables, which contain all available information about past dynamics of diversification (14); this suggests an alternative paradigm for learning about the past from extant timetrees.

## Introduction

A central challenge in evolutionary biology is to explain why some taxonomic groups and some time periods have so many species while others have so few; ultimately this means estimating and explaining variation in rates of speciation and extinction (13). Estimating these rates is crucial to investigating fundamental questions such as the role of biotic and abiotic processes in shaping patterns of species richness (7), how Earth’s biota recover after mass extinction events (15, 16) and whether there are general dynamics that govern how biodiversity accumulates (17). Measuring such rates has taken on a new urgency as we try to understand how anthropogenically induced extinctions compare to “background” rates (18, 19). In the medical domain, “speciation” and extinction rates are key parameters that provide insights into the historical dynamics and future trajectories of viral epidemics (20, 21).

Unfortunately, the vast majority of lineages that have ever lived have not left any trace in the fossil record, and hence we are forced to attempt to reconstruct their diversification dynamics from incomplete data. Indeed, for many groups the fossil record is so incomplete that the primary source of information on past diversification dynamics comes from time-calibrated molecular phylogenies of extant lineages (“extant timetrees”). There is now an abundance of increasingly sophisticated methods for extracting this information, with most state-of-the-art methods fitting variants of a birth-death process (22) to extant timetrees (1–6, 23). These methods have, collectively, been used in thousands of studies and have substantially contributed to our understanding of the drivers of diversity through time. Despite their popularity, there has been a long-lingering doubt about many of these inferences. For one, simulation studies have repeatedly shown that some variables, especially extinction rates, are generally difficult to estimate (11, 24–28). But an even more fundamental issue, which has particularly drawn the attention of paleobiologists, is that there may not be sufficient information in a molecular phylogeny to fully reconstruct historical changes in diversification rates. For example, when speciation and extinction rates vary through time — and there is abundant evidence from the fossil record that they do (13) — mass extinction events can erode much of the signal of preceding diversification dynamics (9, 10, 12), and may themselves even be confused with stagnating speciation rates (29). To date these critical identifiability issues remain poorly understood, and no general theory exists for describing which diversification scenarios can be distinguished from each other and precisely what information on diversification rates is in principle extractable from extant phylogenies.

Here, we present a solution to this problem: We develop a mathematical framework for assessing the identifiability of the general stochastic birth-death process with homogeneous rates, where speciation (“birth”) rates (*λ*) and extinction (“death”) rates (*µ*) can vary over time, and which underlies the majority of existing methods for reconstructing diversification dynamics from phylogenies (5). By considering the full space of possible diversification scenarios (i.e., with arbitrary *λ* and *µ*), rather than special cases (as has been done so far), we reveal a fundamental and surprisingly general property of the birth-death process that has far reaching implications for diversification analyses. Specifically, we show that for any given birth-death model there exists an infinite number of alternative birth-death models that can explain any given extant timetree equally well as the candidate model. This ambiguity persists for arbitrarily large trees and cannot be resolved even with an infinite amount of data using any statistical method. Crucially, these alternative models may appear to be similarly plausible and yet exhibit markedly different features, such as different trends through time in both *λ* and *µ*. Using simulated and real timetrees as examples, we demonstrate how failing to recognize this immense ambiguity may seriously mislead our inferences about past diversification dynamics, shedding doubt on conclusions from countless previous studies. We further show that these sets of “congruent” models can be uniquely identified based on suitably defined composite variables: the “pulled speciation rate”, corresponding to the effective *λ* in the hypothetical absence of extinction and under complete species sampling, or equivalently, the “pulled diversification rate”, corresponding to the effective net diversification rate in the hypothetical case where *λ* is time-independent. Based on either one of these variables, it becomes possible to determine whether different diversification scenarios are at all distinguishable, to explore the full range of plausible scenarios that are consistent with the data, and to make inferences about diversification dynamics without knowing *λ* and *µ* themselves.

### Computing the likelihood of diversification models from lineages-through-time curves

One of the most important features of extant timetrees is the lineages-through-time curve (LTT), which counts the number of lineages at each time in the past that are represented by at least one extant descending species in the tree. The LTT provides a simple visual overview of a tree’s branching density over time and importantly, contains all the information encoded in the tree regarding speciation and extinction rates (30) (see also Supplement S.1.2). This is because the likelihood of a extant timetree under a given birth-death model with homogeneous rates depends solely on the tree’s LTT, but not on any other properties of the tree that do not affect the LTT.

Here we show that an elegant analogous relationship also exists between the likelihood of a tree and the LTT that would be predicted by a given birth-death model. Any given speciation and extinction rates over time, *λ* and *µ*, and the probability that an extant species will be included in the tree *ρ* (present-day “sampling fraction”), can be used to define a “deterministic” diversification process, where the number of lineages through time no longer varies stochastically but according to a set of differential equations (3,31–33; also see Supplement S.1). The LTT predicted by these differential equations (“deterministic LTT”, or dLTT) is a mere theoretical property of the model that resembles the LTT of a tree only if the tree is sufficiently large for stochastic effects to become negligible, and assuming the model is an adequate description of the process that generated the tree. It can be shown, however, that the likelihood of a tree under a given birth-death model can be written purely in terms of the tree’s LTT and the model’s dLTT (Supplement S.1.2). This means that any two models with the same dLTT (conditioned on the number of extant species sampled, *M*_*o*_) yield identical likelihoods for the tree. In the following, we shall therefore call any two models “congruent” if they have the same dLTT for any given *M*_*o*_. Note that two models are either congruent or non-congruent regardless of the particular tree considered, meaning that model congruency is a property of models and not the data (Supplement S.1). Furthermore, the probability distribution of tree sizes generated by a birth-death model, when conditioned on the age of the stem (or crown), is identical among congruent models (Supplement S.1.7). Hence, congruent models have equal probabilities of generating any given timetree and LTT, analogous to how congruent geometric objects exhibit similar geometric properties (discussion in Supplement S.1.8). In the absence of further information or constraints, congruent models cannot possibly be distinguished solely based on extant timetrees, neither through the likelihood nor any other test statistic (such as the *“* statistic (34)). Note that whether or not a given phylogenetic data set is sufficient to statistically distinguish between non-congruent models is an entirely different matter.

### Congruent model sets are infinitely large and infinite-dimensional

The above considerations lead to an important question: For any birth-death model (i.e., with given *λ* and *µ* as functions of time, and a given sampling fraction *ρ*), how many alternative congruent models are there and how could one possibly construct them? To answer this question, we first present an alternative method for recognizing congruent models. Given a number of sampled species *M*_*o*_, a model’s dLTT is fully determined by its relative slope, *λ*_p_ = −*M*^−1^*dM/dτ* (where *M* is the dLTT and *τ* is time before present or “age”). It can be shown that *λ*_p_ is related to the model’s speciation rate as *λ*_p_ = *Pλ*, where *P* (*τ*) is the probability that any lineage that existed at age *τ* survives to the present and is included in the timetree (Supplement S.1.1). In the absence of extinction (*µ* = 0) and under complete species sampling (*ρ* = 1), *λ*_p_ is identical to *λ*, however in the presence of extinction *λ*_p_ is pulled downwards relative to *λ* at older ages, while under incomplete sampling *λ*_p_ is pulled downwards relative to *λ* near the present. We thus henceforth refer to *λ*_p_ as the “pulled speciation rate” of a model. Since a model’s dLTT is fully determined by *λ*_p_ and, reciprocally, *λ*_p_ is fully determined by the dLTT, it becomes evident that two models are congruent if and only if the have the same pulled speciation rate. The latter can also be used to calculate a variable called “pulled diversification rate” (14), defined as:

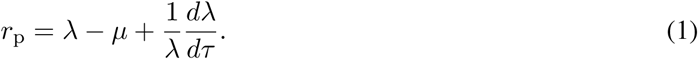

The *r*_p_ is equal to the net diversification rate (*r* = *λ − µ*) whenever *λ* is constant in time (*dλ/dτ* = 0), but differs from *r* when *λ* varies with time. As shown in Supplement S.1.1, *λ*_p_ and *r*_p_ are linked through the following differential equation:

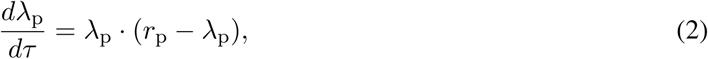

with initial condition *λ*_p_(0) = *ρλ_o_* (where *λ_o_* = *λ*(0) is the present-day speciation rate). Equation (2) reveals that *r*_p_ is completely determined by *λ*_p_ (one can just solve for *r*_p_). Reciprocally, *λ*_p_ is completely determined by *r*_p_ and some initial condition (i.e., *λ*_p_ specified at some fixed time), since one can just solve the differential equation for *λ*_p_ (see solution in Supplement S.1.6). We thus conclude that two birth-death models are congruent if, and only if, they have the same *r*_p_ and the same *λ*_p_ at some time point (for example the same product *ρλ_o_*).

We are now ready to assess the breadth of congruent model sets. Consider a birth-death model with speciation rate *λ >* 0, extinction rate *µ* ≥ 0 and sampling fraction *ρ* ∈ (0, 1]. If we denote *η_o_* = *ρλ_o_*, then for any alternative chosen extinction rate function *µ^*^* 0, and any alternative assumed sampling fraction *ρ^*^* ∈ (0, 1], there exists a speciation rate function *λ^*^ >* 0 such that the alternative model (*λ^*^, µ^*^, ρ^*^*) is congruent to the original model. In other words, regardless of the chosen *µ^*^* and *ρ^*^*, we can find a hypothetical *λ^*^* that satisfies:

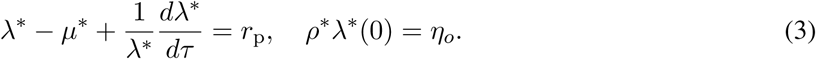

Indeed, to construct such a *λ^*^* one merely needs to solve the following differential equation:

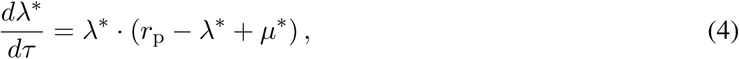

with initial condition *λ^*^*(0) = *η_o_/ρ^*^* (solution given in Supplement S.1.4). The above observation implies that, starting from virtually any birth-death model, we can generate an infinite number of alternative congruent models simply by modifying the extinction rate *µ* and/or the assumed sampling fraction *ρ*. Alternatively, congruent models can be generated by assuming various ratios of extinction over speciation rates, *ϵ* = *µ/λ* (formula in Supplement S.1.5). This set of congruent models — henceforth “congruence class” — is thus infinitely large. The congruence class can have an arbitrary number of dimensions (depending on restrictions imposed a priori on *λ^*^* and *µ^*^*), since *µ^*^* could depend on an arbitrarily high number of free parameters.

For illustration, consider the simulations in Figure 1, showing four markedly distinct and yet congruent birth-death models. The first scenario exhibits a constant *λ* and a temporary spike in *µ* (mass extinction event), the second scenario instead exhibits a constant *µ* and a temporary drop in *λ* (temporary stasis of speciations) around the same time, the third scenario exhibits a mass extinction event at a completely different time and a fluctuating *λ*, while the fourth scenario exhibits an exponentially decaying *µ* and a fluctuating *λ*. These congruent scenarios were obtained simply by assuming alternative extinction rates, and a myriad of other congruent scenarios also exist. Analogous situations can be readily found in the literature. Figures 2A–C, for example, show a birth-death model with exponentially varying speciation and extinction rates, *λ* = *λ_o_e^ατ^* and *µ* = *µ_o_e^βτ^* (where *λ_o_*, *µ_o_*, *α* and *β* are fitted parameters), as commonly considered in other studies (3, 35), fitted to a massive timetree of 79,874 extant seed plant species via maximum likelihood. Simply by modifying the coefficient *β* and choosing *λ* according to Eq. (4), one can obtain an infinite number of congruent and similarly complex scenarios, with even opposite trends over time (Fig. 2B). Similarly, Figs. 2D–F show origination and extinction rates of marine animal genera estimated from fossil data (36), compared to two congruent scenarios, one where the linear trend of *µ* has been reversed (Fig. 2B) and one where *µ* was set to zero (Fig. 2C).

**Figure 1:**
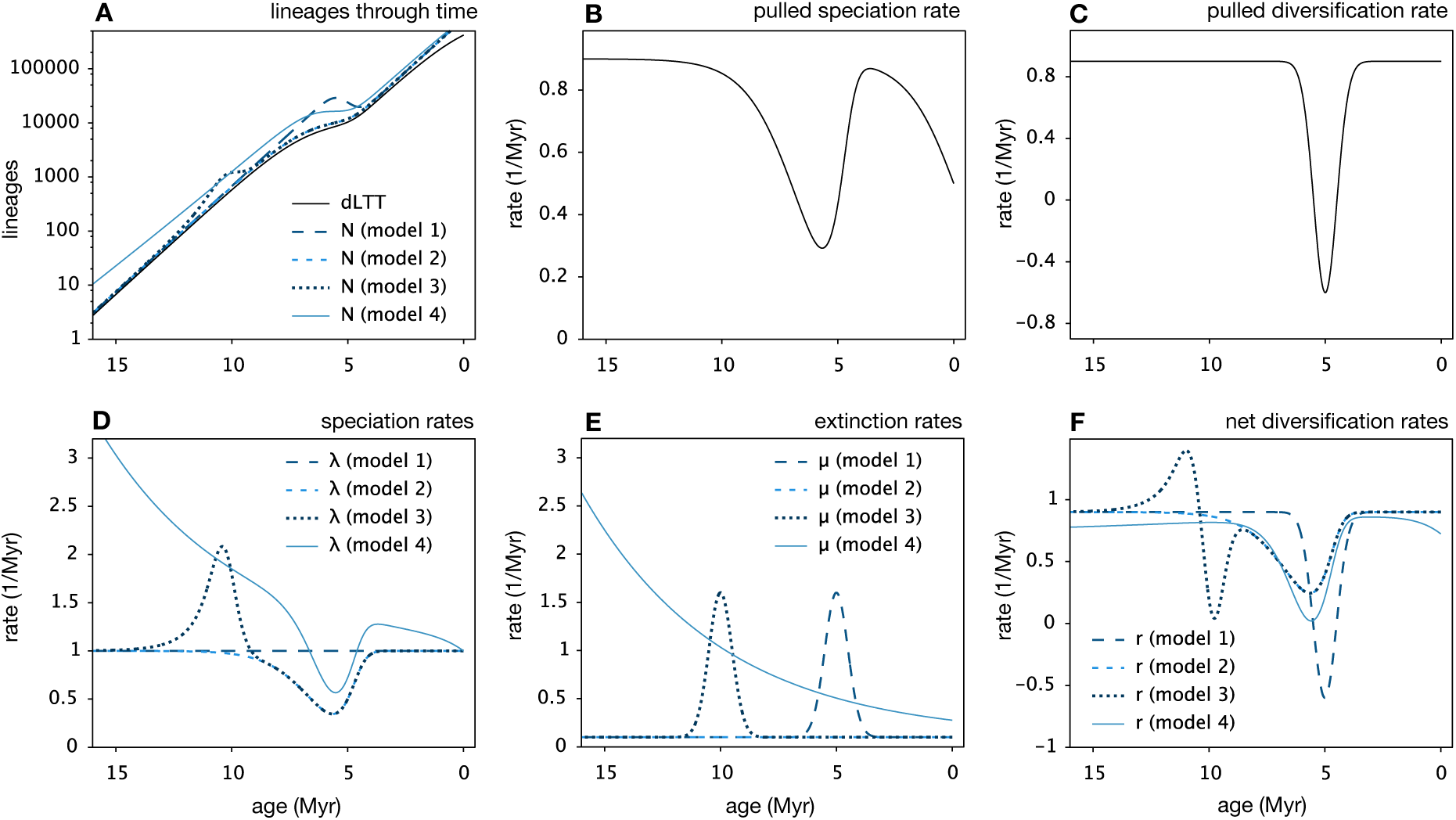
Illustration of congruent birth-death processes (simulations). Example of four hypothetical, congruent yet markedly different birth-death models. The first model exhibits a constant speciation rate and a sudden mass extinction event about 5 Myr before present; the second model exhibits a constant extinction rate and a temporary stagnation of the speciation rate about 5–6 Myr before present; the third model exhibits a mass extinction event about 10 Myr before present and a variable speciation rate; the fourth model exhibits an exponentially decreasing extinction rate and a variable speciation rate. In all models the sampling fraction is *ρ* = 0.5. All models exhibit the same deterministic LTT (dLTT), the same pulled speciation rate (*λ*_p_) and the same pulled diversification rate (*r*_p_), and would yield the same likelihood for any given extant timetree. (A) dLTT and deterministic total diversities (*N*) predicted by the models, plotted over age (time before present). (B) Pulled extinction rate *λ*_p_ of the models. (C) Pulled diversification rate *r*_p_ of the models. (D) Speciation rates (*λ*) of the models. (E) Extinction rates (*µ*) of the models. (F) Net diversification rates (*r* = *λ − µ*) of the models. For additional examples see Supplemental Fig. S1.

**Figure 2:**
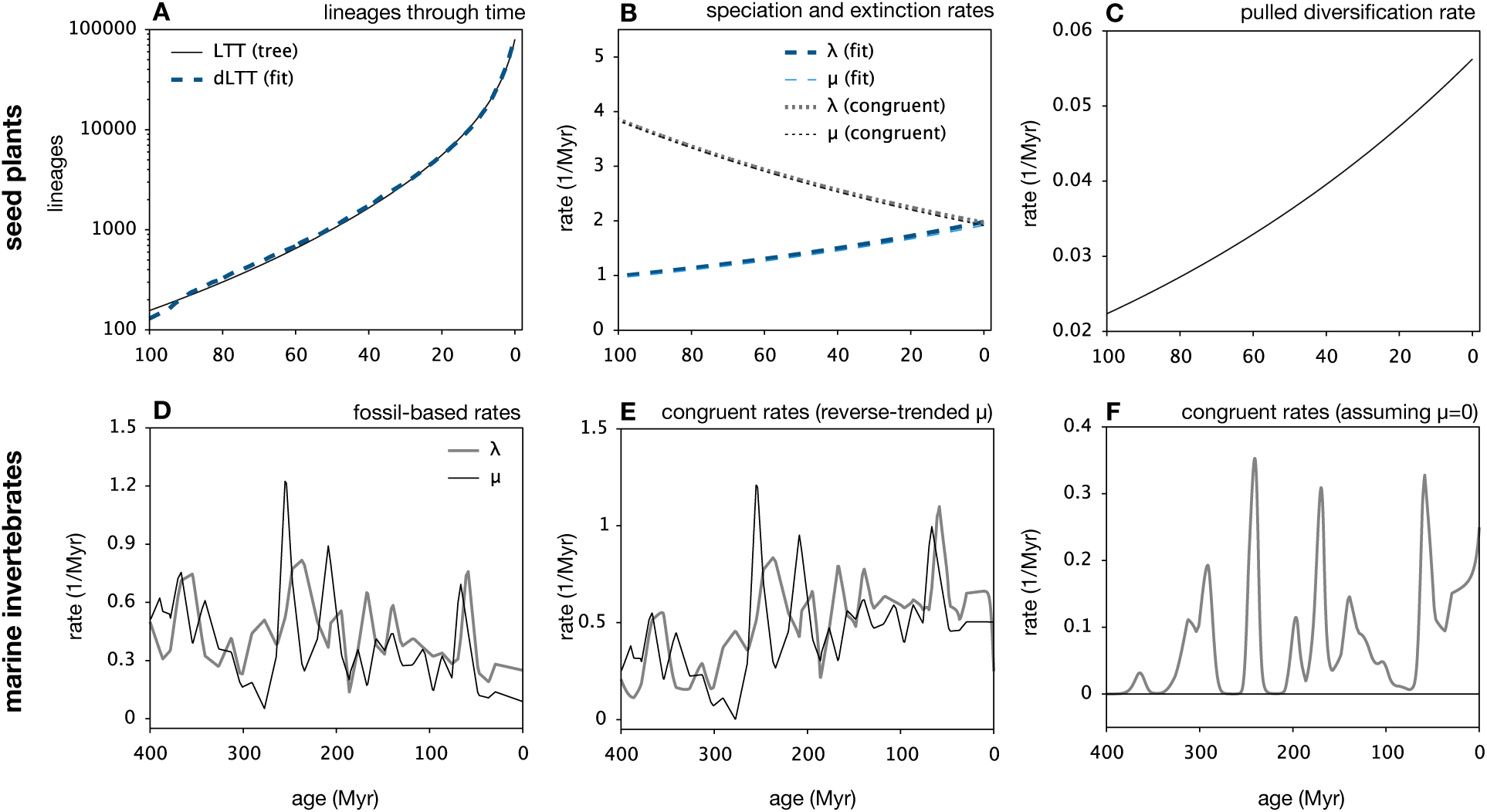
Illustration of congruent birth-death processes (real data). Top row: Birth-death model with exponentially varying *λ* and *µ*, fitted to a reconstruct timetree of 79,874 seed plant species (54) over the past 100 Myr, compared to a congruent model obtained by simply modifying the exponential coefficient of *µ*. (A) LTT of the tree, compared to the dLTT predicted by the two models. (B) Speciation rates (*λ*) and extinction rates (*µ*) of the two models. (C) Pulled diversification rate of the two models. Bottom row: (D) Origination and extinction rates of marine invertebrate genera, estimated from fossil data by Alroy (36). (E) Congruent scenario to D, after reversing the linear trend of *µ*. (F) Congruent scenario to D, assuming zero extinction rate.

Such ambiguities were described previously in special cases. For example, Kubo and Iwasa (31) recognized that a variable *λ* and constant *µ* can be exchanged for a constant *λ* and a variable *µ* to produce the same dLTT; similarly Stadler (29) and Crisp *et al.* (37) observed that simulations of mass extinctions produced similar LTTs as simulations of temporarily stagnating diversification processes. Other previous work on constant-rate birth-death models revealed that alternative combinations of time-independent *λ*, *µ* and *ρ* can yield the same likelihood for a tree (38–41). By generalizing these analyses to the time-variable case, we have revealed that in fact vast expanses of model space are practically indistinguishable.

### Model congruency compromises existing reconstruction methods

Since the likelihood of an extant timetree can be expressed purely in terms of *r*_p_ and the product *ρλ_o_* (Supplement S.1.6), or alternatively purely in terms of *λ*_p_ (Supplement S.1.3), extant timetrees only provide information about the congruence class of a generating process and not the actual speciation and extinction rates. This identifiability issue can be interpreted as follows: Since all information available on past diversification dynamics (representable by birth-death models, to be precise) is encoded in a single curve, namely the LTT, one should not expect to be able to “extract” from it two independent curves (*λ* and *µ*) without additional information, as this would essentially double the amount of information at hand.

In order to estimate *λ* and *µ*, previous phylogenetic studies have been imposing strong and largely arbitrary constraints. For example, many studies assume that *λ* or *µ* vary exponentially through time (42). However this specific functional form is rarely justified biologically, and alternative functional forms of comparable simplicity and shape (e.g., the logistic function, or the Gauss error function) can be envisioned. Normally one expects that, regardless of which of these functional forms is considered, with sufficient data fitting any of these forms will lead to qualitatively similar trends and shapes. This expectation simply does not hold in our case, because the best-fitting representative within any given model set will generally only be the one closest to the congruence class of the true process, rather than closest to the true process itself (Fig. 3). Consequently, fitting alternative functional forms (i.e., alternative model sets) can result in drastically different inferences with alternative trends in *λ* and *µ* over time, even if the each functional form used is in principle adequate for approximating the true historical *λ* and *µ* (examples in Supplement S.6 and Supplemental Fig. S5). This conclusion applies to virtually any model set, including birth-death shift models where *λ* and *µ* change at discrete time points (43).

**Figure 3:**
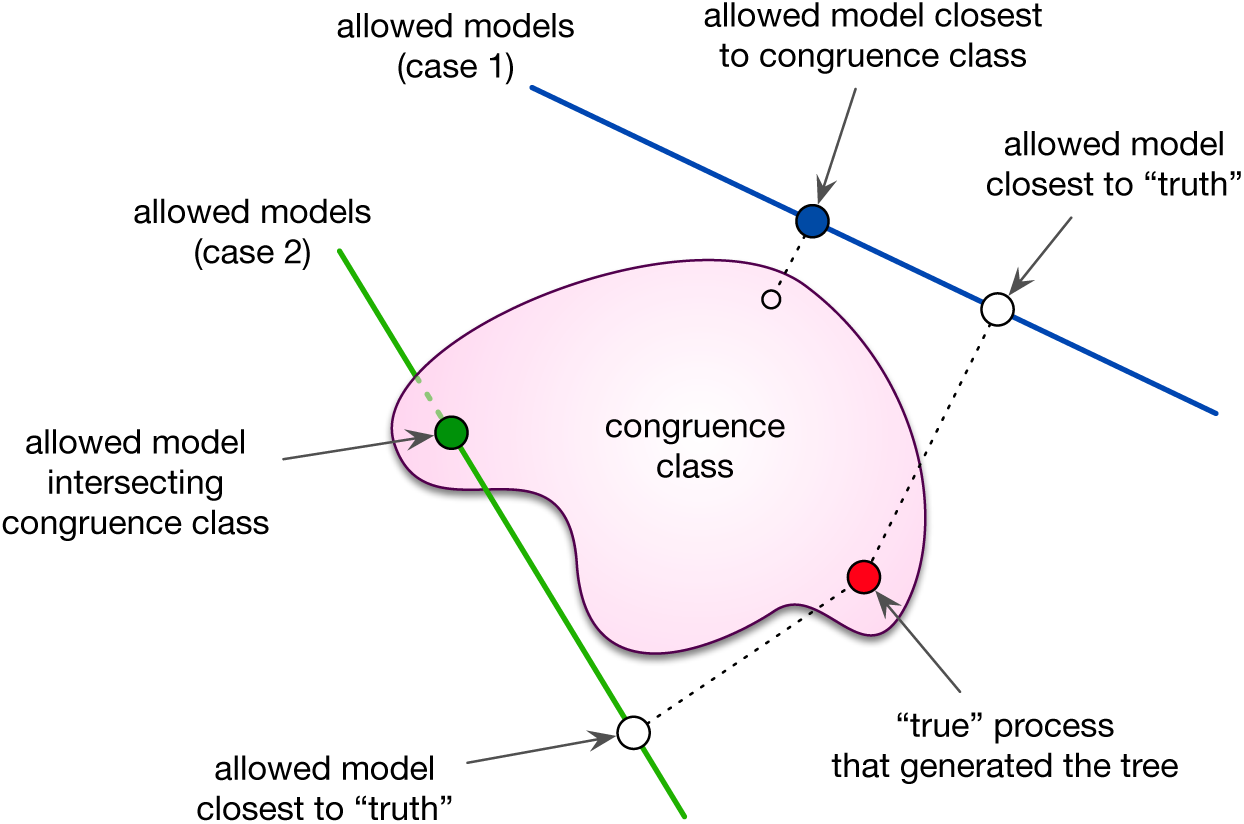
Conceptual implications for reconstructing diversification history. Conceptual illustration of the limited identifiability of a diversification process, assumed to be adequately described by some unknown birth-death model (red circle, henceforth “true process”). The congruence class of the true process is shown as a sub-space comprising a continuum of alternative models (pink area). In practice, maximum-likelihood model selection is performed among a parameterized low-dimensional set of allowed models, the precise nature of which can vary from case to case, for example depending on assumed functional forms for *λ* and *µ* or the number of allowed rate shifts (43). The two continuous lines shown here represent two alternative cases of allowed model sets (e.g., considered in two alternative studies), from within each the model closest to the truth is (ideally) sought. In each case, however, likelihood-based model selection will converge towards the allowed model closest to the congruence class (blue and green filled circles), which in general is not the allowed model actually closest to the true process (white circles). This identifiability issue persists even for infinitely large datasets.

We stress that common model selection methods based on parsimony or “Occam’s razor”, such as the Akaike Information Criterion (AIC; 44) and the Bayesian Information Criterion (BIC; 45) that penalize excessive parameters, generally cannot resolve this issue for multiple reasons. First, these information criteria were designed to minimize the error of future model-based predictions by avoiding overfitting to finite data, and not to identify the actual process that generated the data — these are very different scientific goals with well-known trade-offs. There is no reason to believe that the simplest scenario in a congruence class will be the one closest to the truth. Indeed, even if the true model is included in a congruence class, it will almost always be the case that there are both simpler and more complex scenarios within the same congruence class (e.g., Figs. 2D–F) and, crucially, all of these alternative models remain equally likely even with infinitely large datasets. Second, if one were to apply AIC or BIC, it is unclear how to quantify the complexity of a diversification scenario in comparison with alternative scenarios, which may be described using different functional forms. It is tempting to think that one could simply count the number of parameters. However, any given curve can be written using various alternative functional forms parameterized in distinct ways (recall that ultimately we wish to approximately estimate the curves *λ* and *µ*, not the parameters of some functional form); the number of parameters is a property of parameterized sets of curves, not of a single curve. Even if that were not the case, the number of parameters conventionally associated with a given functional form need not necessarily reflect our intuition about complexity: Is a linear extinction rate (*µ* = *α* + *β ⋅ τ*, two parameters) more or less complex than an exponentially decaying rate (*µ* = *αe^βτ^*) or an oscillating rate of the form *µ* = *α* sin^2^(*βτ*)? In addition, different members of a congruence class may be described with different functional forms involving the same number of parameters. For example, the diversification scenario with linear extinction rate (*µ* = *α* + *β ⋅ τ*) and constant speciation rate (*λ* = *“*) (3 parameters, assuming complete species sampling) is congruent to an alternative and markedly different scenario with zero extinction rate (*µ^*^* = 0) and *λ^*^* defined as the solution to the differential equation *dλ^*^/dτ* = *λ^*^* ⋅ (*γ − α − βτ − λ*) with initial condition *λ^*^*(0) = *γ* (again 3 parameters); there is no reason to prefer one congruent scenario over the other based on the number of parameters or biological realism. Third, even when fitting models of the same functional form, as explained above the maximum-likelihood model will a priori tend to be the one closest to the congruence class of the true process, rather than the true process itself, and neither AIC nor BIC would resolve this (since all other allowed models would have the same number of parameters but lower likelihood). Supplemental Fig. S6 shows examples where maximum-likelihood fitted models, chosen among a wide range of model complexities based on AIC, grossly fail to estimate the true rates even when fitting to a massive tree with 1,000,000 tips, despite the fact that the models could in principle have accurately captured the true rates.

Previous studies have not recognized the breadth of this issue because they typically only consider a limited set of candidate models at a time, both when analyzing real datasets as well as when assessing parameter identifiability via simulations; as a result, previous studies have been (un)lucky enough to not compare two models in the same congruence class (see Supplements S.2 and S.3 for reasoning). For example, if a tree was generated by an exponentially decaying *λ* and *µ* (e.g., via simulations), then fitting an exponential functional form will of course yield accurate estimates of the exponents; however if the generating process was only approximately exponential and better described by another gradually decaying function, then fitting an exponential curve could even lead to opposite trends (examples in Fig. 2 and Supplement S.6).

It is important to realize that congruent scenarios can have markedly different macroevolutionary implications. For example, Steeman *et al.* (46) reconstructed past speciation rates of Cetaceans (whales, dolphins, and porpoises) based on an extant timetree and using maximum-likelihood (assuming *µ* = 0). Steeman *et al.* (46) found a temporary increase of *λ* during the late Miocene-early Pliocene (Fig. 4), suggesting a potential link between Cetacean radiations and concurrent paleoceanographic changes. However, alternatively to assuming *µ* = 0, one could assume that *µ* was close to *λ*, consistent with common observations from the fossil record (13). For example, by setting *µ* = 0.9 · *λ* one obtains a congruent scenario in which *λ* no longer peaks during the late Miocene-early Pliocene but instead exhibits a gradual slowdown throughout most of Cetacean evolution (Fig. 4B). Both scenarios are similarly complex and both could have generated the timetree at equal probabilities. Even the common methodological decision to estimating net diversification (*r* = *λ* − *µ*) rather than *λ* and *µ* separately (8, 11), is no longer meaningful in light of our results; the shape of *r* is not conserved across a congruence class. Likewise, the models in a congruence class will not share “average” (however defined) rates; hence absolute rate estimates (1), which have been used to estimate broad macroevolutionary patterns (17) and background rates of extinction (19), are also unlikely to be accurately reconstructed. These issues are likely also present in more complex models with additional free parameters, for example where some clades exhibit distinct diversification regimes (42, 47). In such situations, the tree can always be decomposed into a set of sub-trees with distinct LTT curves, each of which is subject to the same identifiability issues as described here. Our findings thus shed doubts over a lot of previous work on diversification dynamics, including some of the conclusions from our own work (14, 17).

**Figure 4:**
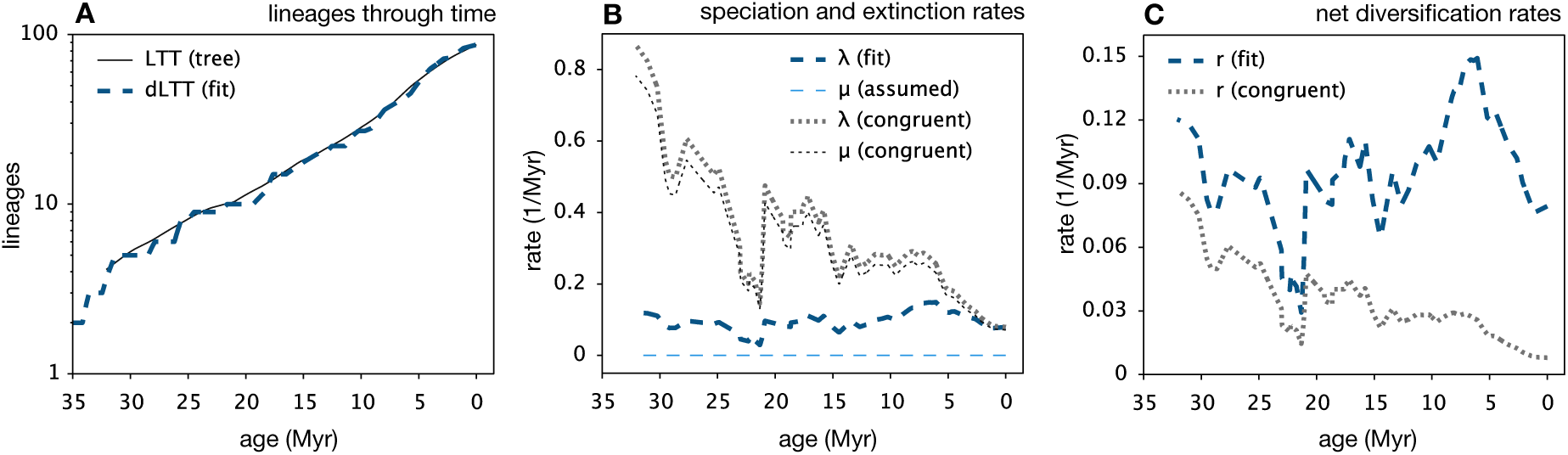
Previous studies have likely over-interpreted phylogenetic data. Time-dependent birth-death model fitted to a nearly-complete Cetacean timetree by Steeman *et al.* (46) under the assumption of zero extinction rates (*µ* = 0), compared to a congruent model where the extinction rate is close to the speciation rate (*µ* = 0.9*λ*). (A) LTT of the tree, compared to the dLTT predicted by the two models. (B) Speciation rates (*λ*) and extinction rates (*µ*) of the two models. (C) Net diversification rates (*r* = *λ − µ*) of the two models. The original fitted rates were used by Steeman *et al.* (46) to link Cetacean diversification dynamics to past paleoceanographic changes.

### Ways forward

Our findings for birth-death models of diversification are closely analogous to classic results from coalescent theory in population genetics (48, 49), where an infinite number of models can give rise to the same drift process as the idealized Wright-Fisher model. This realization was profoundly important for the field; it focused researchers’ attention on the dynamics of the effective population size *N*_*e*_, a composite but identifiable parameter that is the same for all models with the same Wright-Fisher drift process, rather than on the actual (but non-identifiable) historical demography, sex ratios etc. of the population. Consequently, the field has adapted and developed a plethora of tools to infer changes in *N*_*e*_ through time and across the genome. Here we have found an analogous generality, and have confirmed previous suspicions that many historical diversification scenarios may not be distinguishable using extant phylogenies alone (10, 13, 30, 31). We have shown that such congruent scenarios can be defined in terms of the *λ*_p_, or equivalently, in terms of the *r*_p_ and *ρλ*_*o*_, all of which are identifiable provided sufficient data (indeed, for sufficiently large trees these variables can be directly calculated from the slope and curvature of the LTT; 14). Each congruence class contains exactly one model with *µ* = 0 and *ρ* = 1, which is also the only model where *λ* = *λ*_p_; hence the pulled speciation rate can be interpreted as the effective speciation rate generating the congruence class’s dLTT in the absence of extinctions and under complete species sampling. Similarly, each congruence class contains an infinite number of models with time-independent *λ*, and for these models *r*_p_ = *r*; hence the pulled diversification rate can be interpreted as the effective net diversification rate if *λ* was time-independent. It is in this way that *λ*_p_ and *r*_p_, being identifiable and “effective” rates in idealized scenarios, are analogous to *N*_*e*_ in population genetics.

Of course fossil data could in principle help resolve the issues highlighted here, for example via fossilized-birth-death models (50, 51) or birth-death-chronospecies models (52), however for a large number of taxa (e.g., all prokaryotes and many soft-bodied eukaryotes) fossil data are virtually non-existent. Rather than attempting to estimate *λ* and *µ*, one can instead estimate *λ*_p_, *r*_p_ and *ρλ*_*o*_ (and *λ*_*o*_ if *ρ* is known) either using likelihood methods (Supplement S.5) or based on the slope and curvature of a tree’s LTT (14). Our previous work(14) has shown that *r*_p_ itself can yield valuable insight into diversification dynamics and can be useful for testing alternative hypotheses (also see Supplement S.4). Using simulations, for example, we found that sudden rate transitions, for example due to mass extinction events, usually lead to detectable fluctuations in *r*_p_; therefore, a relatively constant *r*_p_ over time would be indicative of constant — or only slowly changing — speciation and extinction rates (14). One can also obtain other useful composite variables from *λ*_p_, *r*_p_ and *ρλ*_*o*_. For example, in cases where *ρ* is known one can obtain the “pulled extinction rate”, defined as *µ*_p_:= *λ*_*o*_ − *r*_p_ (14). Note that *µ*_p_(*τ*) is equal to the extinction rate *µ*(*τ*) if *λ* has been constant from *τ* to the present, but differs from *µ* in most other cases. The present-day *µ*_p_ is related to the present-day *µ* as follows:

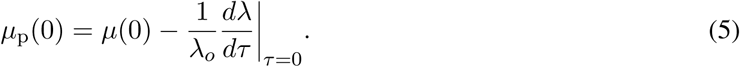

Hence if the present-day speciation rate changes only slowly, the present-day *µ*_p_ will resemble the presentday *µ*. Further, since *µ*(0) is non-negative, we can obtain the following one-sided bound for the rate at which *λ* changes at present:

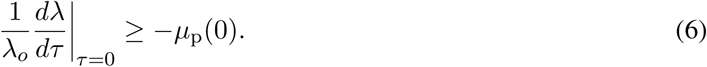

If a macroevolutionary question is only concerned with recent speciation events (7) then one can test hypotheses using *λ*_*o*_, which can be readily identified if *ρ* is known.

## Conclusions

We have shown that for virtually any candidate birth-death process, suspected of having generated some extant timetree, there exists an infinite number of alternative and markedly different birth-death processes that could have generated the timetree with the same likelihood. Without further information or prior constraints on plausible diversification scenarios, extant timetrees alone cannot be used to reliably infer speciation rates (except at present-day), extinction rates or net diversification rates, raising serious doubts over a multitude of previous estimates of past diversification dynamics. Our work could thus explain why frequently diversification dynamics observed in the fossil record are in great disagreement with phylogenetics-based inferences (3, 5, 10, 13, 46).

On a more positive note, we resolved a long-standing debate and precisely clarified what information can indeed be extracted from extant timetrees alone — namely *λ*_p_, *r*_p_, the product *ρλ*_*o*_ (and *λ*_*o*_ if *ρ* is known), and any other variables that can be expressed in terms of *λ*_p_, *r*_p_ and *ρλ*_*o*_. These identifiable variables not only tell us when two models are in principle distinguishable but they can themselves yield valuable insight into past diversification dynamics. We see these as analogous to the concept of effective population size in population genetics — we cannot uniquely determine the exact sequence of events that led to current diversity, but by blending out some of the historical details we could potentially gain powerful and robust insights into general macroevolutionary phenomena.

## Code availability

Computational methods used for this article, including functions for simulating birth-death models, for constructing models within a given congruence class, for calculating the likelihood of a congruence class, and for directly fitting congruence classes (either in terms of *λ*_p_ or in terms of *r*_p_ and *ρλ*_*o*_) to extant timetrees (Supplement S.5), are implemented in the R package castor (53).

### Acknowledgments

S.L. was supported by a startup grant by the University of Oregon, USA. M.W.P. was supported by an NSERC Discovery Grant. We thank L. Harmon, S. Otto, A. MacPherson, D. Schluter, T.J. Davies, M. Whitlock, L.F. Henao Diaz, K. Kaur, J. Uyeda, D. Caetano, J. Rolland, L. Parfrey, and A. Mooers for insightful comments on this work.

## Author contributions

Both authors conceived the project and contributed to the writing of the manuscript. S.L. performed the mathematical calculations and computational analyses.

## Supplemental Information

### S.1 Mathematical derivations

In the following, we provide mathematical derivations for various claims made in the main article. Some parts can be found in previous literature (1, 2, 3, 4, 5, 6), but are included here for completeness.

### S.1.1 General considerations

We begin with listing some basic mathematical properties of deterministic birth-death models that will be of use at various later stages. Our starting point is some time-dependent speciation rate *λ*, some time-dependent extinction rate *µ* and some sampling fraction *ρ* (fraction of extant species included in the tree). Let *τ* denote time before present (“age”). The deterministic total diversity, i.e. the number of species predicted at any point in time according to the deterministic model, and conditional upon *M*_*o*_ extant species having been sampled at present-day, is obtained by solving the following differential equation backward in time:

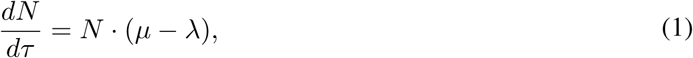

with initial condition *N* (0) = *M*_*o*_/*ρ*, i.e.:

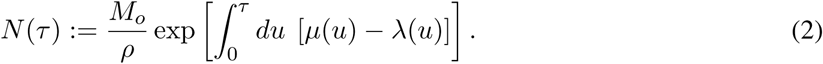

The deterministic LTT (dLTT), i.e. the number of lineages represented in the final extant timetree at any time point according to the deterministic model, is given by:

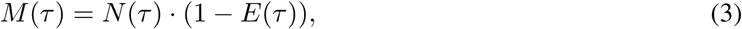

where *E*(*τ*) is the probability that a lineage extant at age *τ* will be missing from the timetree (either due to extinction or not having been sampled). As explained by Morlon *et al.* (5), the extinction probability *E* satisfies the differential equation:

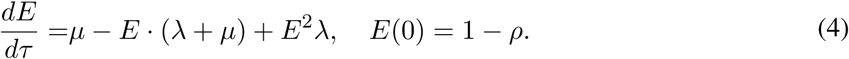

We mention that the solution to Eq. (4) is provided by Morlon *et al.* (5, Eq. 2). Taking the derivative of both sides in Eq. (3), and then using Eq. (4) to replace *dE/dτ* as well as Eq. (1) to replace *dN/dτ* quickly leads to the differential equation:

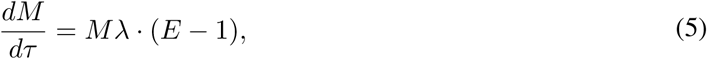

with initial condition *M* (0) = *M*_*o*_. The solution to this differential equation is:

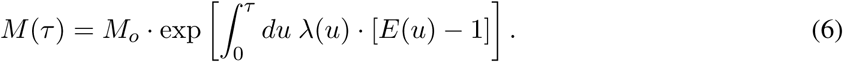

Observe that *E* is a property purely of the model, and does not depend on the particular tree considered; together with Eq. (6), this shows that any two models either have equal dLTTs for any given tree or they have non-equal dLTTs for any given tree. Hence, model congruency is a property of two models, regardless of tree.

Defining the relative slope of the dLTT:

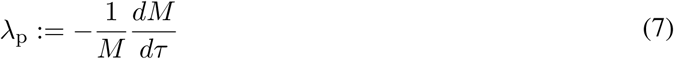

allows us to write Eq. (5) as follows:

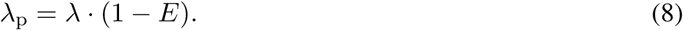

We note that *P* (*τ*):= 1 − *E*(*τ*) is the probability that a lineage extant at age *τ* is represented in the extant timetree. *P* can thus be interpreted as a generalization of the present-day sampling fraction *ρ* to previous times. In fact, trimming a timetree at some age *τ*_*o*_ > 0 (i.e., omitting anything younger than *τ*_*o*_) would yield a new (shorter) timetree, whose tips are a random subset of the lineages that existed at age *τ*_1_, each included at probability *P* (*τ*_*o*_).

As becomes clear in Eq. (8), in the absence of extinction and if *ρ* = 1, the relative slope *λ*_p_ becomes equal to the speciation rate *λ*; in the presence of extinction *λ*_p_ is artificially pulled downwards relative to *λ* towards the past. Reciprocally, under incomplete sampling *λ* is artificially pulled downwards near the present. We shall therefore henceforth call *λ*_p_ the “pulled speciation rate”.

Taking the derivative on both sides of Eq. (8) and using Eq. (4) to replace *dE/dτ* leads to:

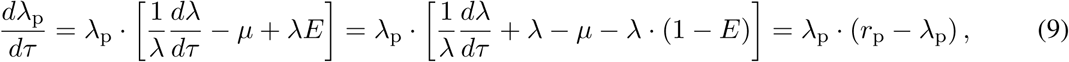

where we defined the “pulled diversification rate”:

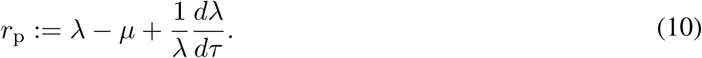

Rearranging terms in Eq. (9) yields:

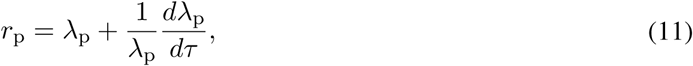

which shows that *r*_p_ can be directly calculated from the dLTT.

#### S.1.2 The likelihood in terms of the LTT and dLTT

In the following we show how the likelihood of an extant timetree under a birth-death model can be expressed purely in terms of the tree’s LTT and the model’s dLTT. We begin with the case where the stem age is known and the likelihood is conditioned on the survival of the stem lineage; the alternative case where only the crown age is known is very similar and will be discussed at the end.

Our starting point is the likelihood formula described by Morlon *et al.* (5):

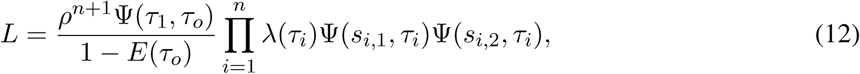

where *n* is the number of branching points (internal nodes), *τ*_*o*_ is the age of the stem, *τ*_1_ > *τ*_2_ > .. > *τ*_*n*_ are the ages (time before present) of the branching points, *s*_*i*,1_, *s*_*i*,2_ are the ages at which the daughter lineages originating at age *τ*_*i*_ themselves branch (or end at a tip), *ρ* is the tree’s sampling fraction (fraction of present-day extant species included in the tree), *E*(*τ*) is the probability that a single lineage that existed at age *τ* would survive to the present and be represented in the tree (5, Eq. 2 therein), *Ψ* is defined as:

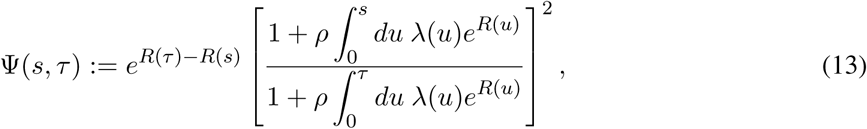

and *R*(*τ*) is defined as:

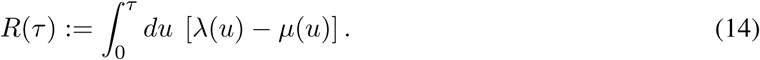

It is straightforward to confirm that *Ψ* satisfies the property *Ψ*(*s, τ*) = *Ψ*(0, *τ*)/*Ψ*(0, *s*); using this property in Eq. (12) leads to:

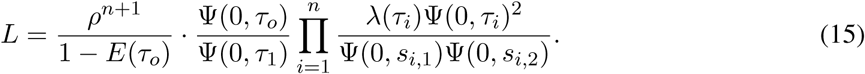

Since each internal node except for the root is the child of another internal node, the enumerator and denominator in Eq. (15) partly cancel out, eventually leading to:

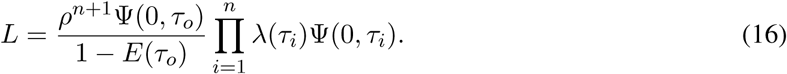

Since the set of branching times *τ*_*i*_ is completely determined by the LTT (branching events correspond to jumps in the LTT), we conclude that the likelihood of a tree is entirely determined by its LTT.

Further, from Eq. (11) we know that the model’s dLTT satisfies:

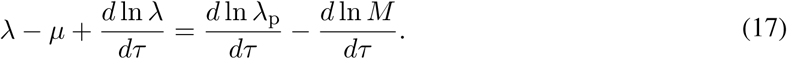

Integrating both sides of Eq. (17) yields:

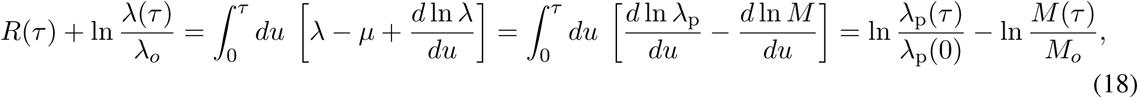

where *M*_*o*_ is the number of extant species included in the timetree. Hence:

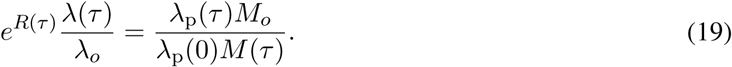

Using Eq. (19) in Eq. (13) yields:

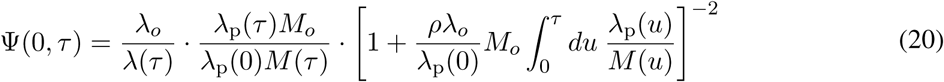

Recall that *ρλ*_*o*_ = *λ*_p_(0) according to Eq. (8), so that Eq. (20) can be written as:

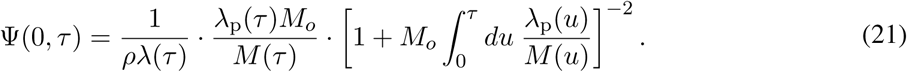

Note that:

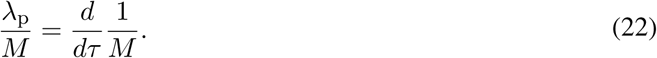

Hence, Eq. (21) can be further simplified to:

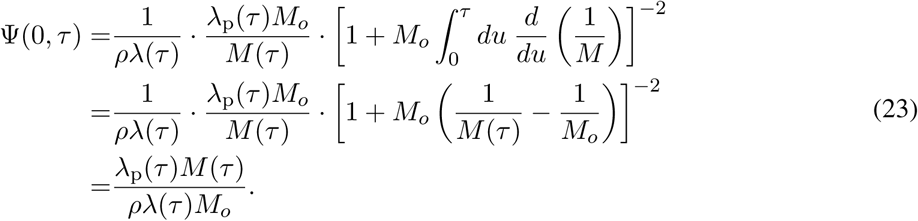

Inserting Eq. (23) into the likelihood formula (16) yields:

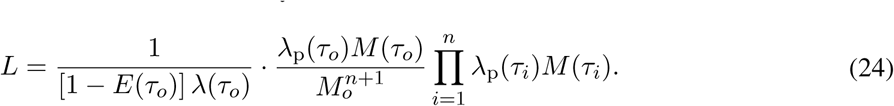

Recall that (1 − *E*) *λ* = *λ*_p_ according to Eq. (8), which when inserted into (24) yields:

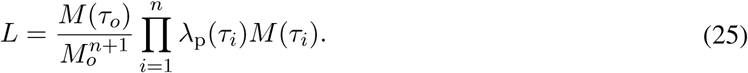

Since *λ*_p_*M* = *−dM/dτ*, Eq. (25) becomes:

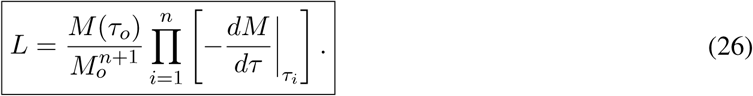

A corollary of Eq. (26) is that for any given extant timetree, any two models with the same dLTT will also any yield the same likelihood.

Note that the likelihood in Eq. (12) or equivalently Eq. (26) is conditioned upon the survival of the stem lineage, assuming that the stem age is known. If the stem age is unknown the likelihood should be conditioned upon the splitting at the root and the survival of the root’s two daughter-lineages, as follows:

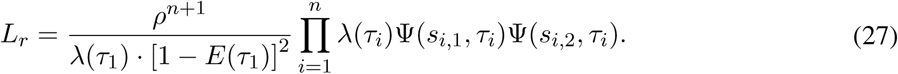

Note that Eq. (27) can be obtained from (12) by setting the stem age equal to the crown age (*τ*_*o*_ = *τ*_1_) and adjusting the conditioning. Following a similar procedure as above, it is easy to show that *L*_*r*_ can be expressed in the following alternative forms:

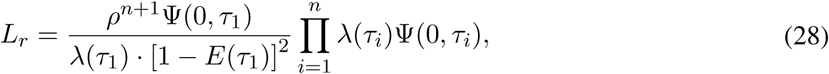

and

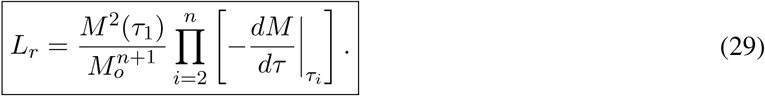

#### S.1.3 The likelihood in terms of *λ*_p_

In the following we show how the likelihood of an extant timetree under a birth-death model can be expressed purely in terms of the tree’s LTT and the model’s pulled speciation rate *λ*_p_.

We begin with the case where the stem age is known and the likelihood is conditioned on the survival of the stem lineage. Our starting point is the likelihood formula in Eq. (26):

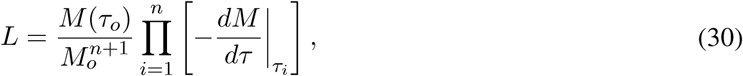

where *M* is the dLTT and *M*_*o*_:= *M* (0). From Eq. (7) it is easy to obtain the following relationship between *M* and *λ*_p_:

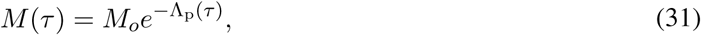

where we defined:

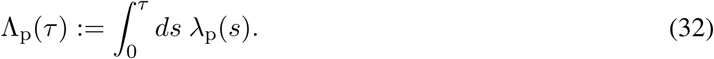

Inserting Eq. (31) into Eq. (30) yields:

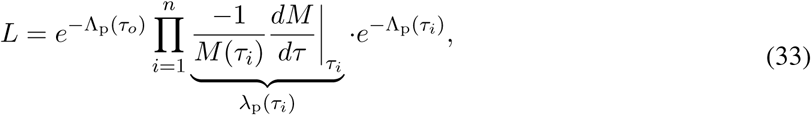

and hence:

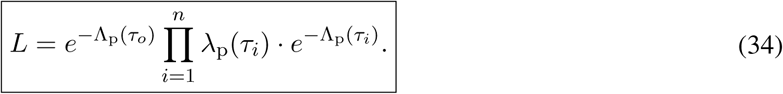

If only the crown age is known and the likelihood is conditioned on the splitting at the root and the survival of the root’s two daughter-lineages (likelihood formula in Eq. (29)), we instead obtain the expression:

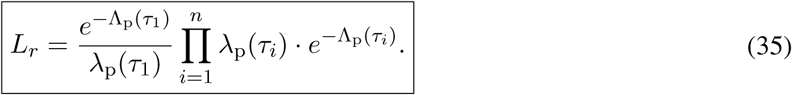

### S.1.4 Calculating *λ* from *r*_p_ and *µ*

In the following we provide the general solution to the differential equation (4) in the main article:

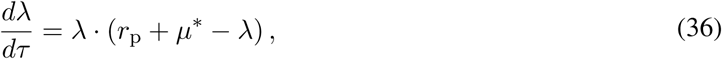

with initial condition:

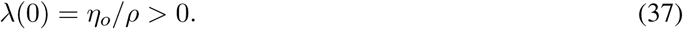

We assume that *r*_p_ and *µ*\* are sufficiently “well-behaved”, specifically that they are integrable over any finite interval. Observe that Eq. (36) is an example of a Bernoulli-type differential equation, as it can be written in the standard form:

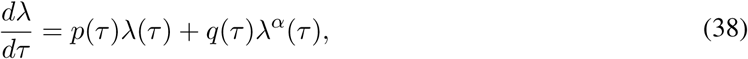

where *α* = 2, *p* = *r*_p_ + *µ*\* and *q* = −1. Using the standard technique for solving Bernoulli differential equations (i.e., substituting *u* = *λ*^1−*α*^ to obtain a linear differential equation for *u*), it is straightforward to obtain the solution:

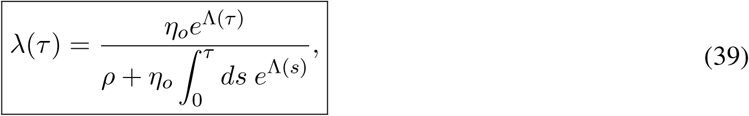

where we defined:

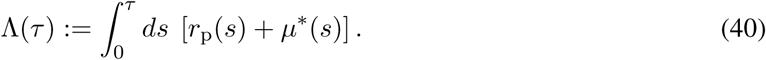

Note that the solution in Eq. (39) is strictly positive and continuous, and hence *λ* is indeed a valid speciation rate.

For future reference, we mention that the above solution can be easily generalized to the case where the “initial condition” for *λ* is given at some arbitrary age *τ*_1_, rather than at present-day. Specifically, the solution to the differential equation:

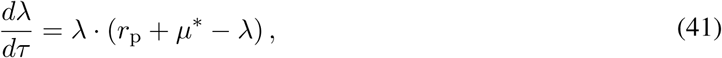

with condition:

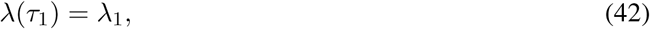

is given by:

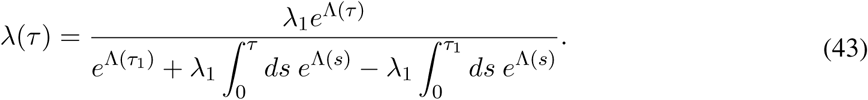

#### Special cases

- In the special case where *r*_p_ and *µ*\* are time-independent and *r*_p_ + *µ*\* ≠ 0, the solution in Eq. (39) takes the form:

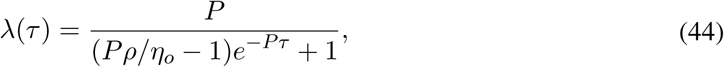

where *P* = *r*_p_ + *µ*\*.
- If and only if *µ*\*(*τ*) = *η*_o_/*ρ* − *r*_p_(*τ*), the solution in Eq. (39) is time-independent:

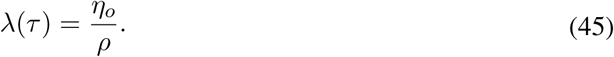 Hence, for a fixed *ρ*, a congruence class can include at most one model with constant speciation rate; it includes exactly one model with constant speciation rate if and only if *η*_*o*_/*ρ* ≥ max_*τ*_ *r*_p_(*τ*).

##### S.1.5. Calculating *λ* from *r*_p_ and *ε*

In the following we show how the speciation rate *λ* can be calculated from the pulled diversification rate *r*_p_, the present-day speciation rate *λ*_*o*_ and the ratio of extinction over speciation rate, *ε*:= *µ/λ*. Specifically, we provide the general solution to the following differential equation:

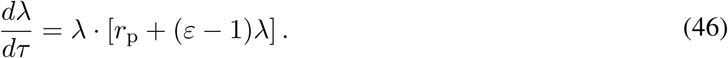

We assume that *r*_p_ and *ε* are sufficiently “well-behaved”, specifically that they are integrable over any finite interval. Observe that Eq. (46) is an example of a Bernoulli-type differential equation, as it can be written in the standard form:

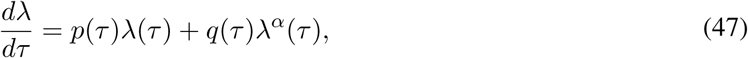

where *α* = 2, *p* = *r*_p_ and *q* = *ε* − 1. Using the standard technique for solving Bernoulli differential equations (i.e., substituting *u* = *λ*^1−*α*^ to obtain a linear differential equation for *u*), it is straightforward to obtain the solution:

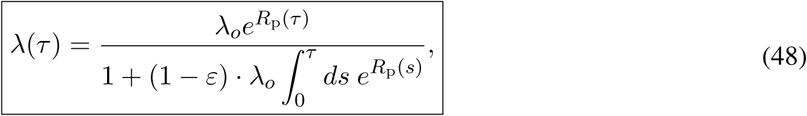

where we defined:

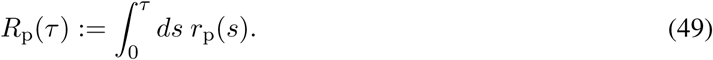

In the special case where *r*_p_ is time-independent and non-zero, the solution in Eq. (48) simplifies to:

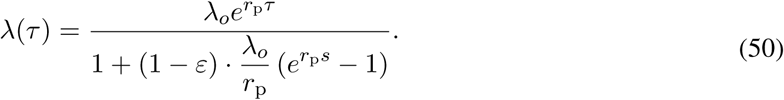

##### S.1.6 The likelihood in terms of the *r*_p_

In the following we show how the likelihood of a tree under a birth-death model can be expressed solely in terms of the model’s pulled diversification rate *r*_p_ and the product *ρλ*_*o*_. We first consider the case where the stem age is known and the likelihood is conditioned on the survival of the stem lineage (5); the alternative case where only the crown age is known and the likelihood is conditioned upon the survival of the root’s two daughter lineages (Eq. 28) can be treated similarly and is briefly mentioned at the end.

Our starting point is the likelihood formula in Eq. (16), Supplement S.1.2. Define:

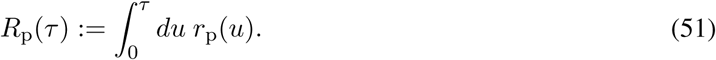

Then from the definition of *r*_p_ (Eq. 1 in the main article) we have:

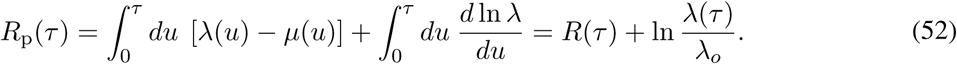

Exponentiating (52) and rearranging yields:

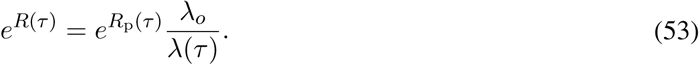

Inserting Eq. (53) into the definition of Ψ in Eq. (13) yields:

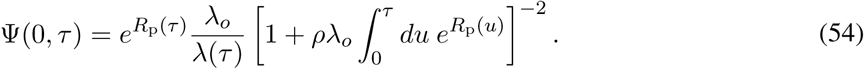

Inserting Eq. (54) into the likelihood formula (16) yields:

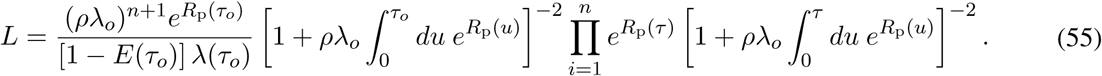

Recall that (1 − *E*)*λ* = *λ*_p_ according to Eq. (8), which when inserted into Eq. (55) yields:

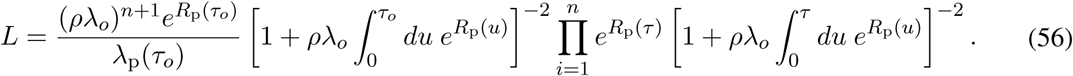

From Eqs. (8) and (11) we know that *λ*_p_ satisfies the initial value problem (Bernoulli differential equation):

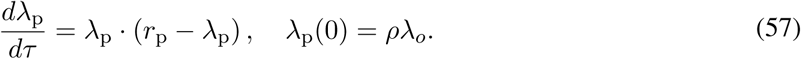

It is straightforward to verify that the solution to Eq. (57) is given by:

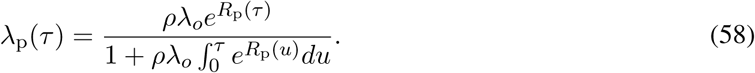

Inserting the solution (58) into Eq. (56) yields the following expression for the likelihood:

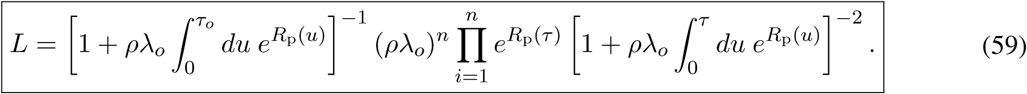

In the alternative case where only the crown age is known, and the likelihood is conditioned on the splitting at the root and the survival of the root’s two daughter lineages, we obtain the following expression for the likelihood:

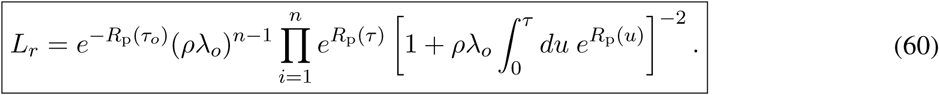

##### S.1.7 Congruent models have the same probability distribution of generated tree sizes

In the following, we show that the distribution of extant timetree sizes generated by a birth-death model, either conditional upon the age and survival of the stem, or conditional upon the age of the root and the survival of its two daughter lineages, is the same for all models in a congruence class.

Consider a birth-death process with parameters (*λ, µ, ρ*), starting from a single lineage at some time before present *τ*_*o*_ and ultimately resulting in a timetree at age 0, comprising only extant species that are included at some probability *ρ*. The probability that the timetree will comprise *n* tips can be expressed using formulas first derived by Kendall *et al.* (7):

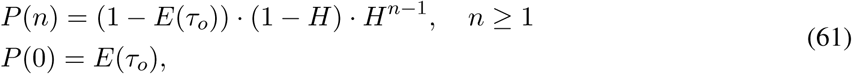

where *E*(*τ*_*o*_) is the probability that a lineage existing at age *τ*_*o*_ will be missing from the timetree (as defined previously), *H* is defined as:

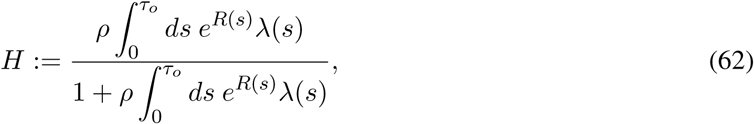

and *R* was previously defined in Eq. (14). Note that the formula in Eq. (61) can be readily obtained using equations 8, 10b and 11 in (7), after setting the time variable therein equal to *τ*_*o*_ (i.e. *t* = *τ*_*o*_), switching from time to age (*τ* = *τ_o_ − t*), and adding the term *−δ*(*τ*) ln *ρ* to the extinction rate (where *δ* is the Dirac distribution, peaking at age 0) to account for incomplete species sampling. As shown previously in Eq. (53), we have

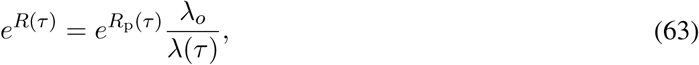

where *R*_p_ is defined as:

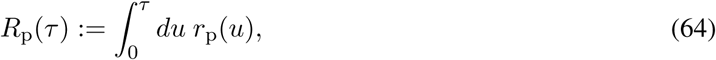

and *r*_p_ is the pulled diversification rate. Inserting Eq. (63) into Eq. (62) allows us to write *H* as follows:

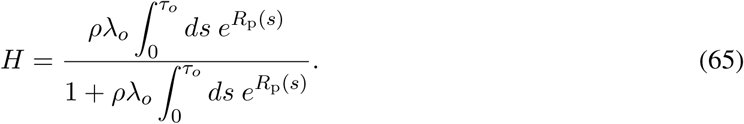

Since *ρλ*_*o*_, *r*_p_ and *R*_p_ are the same for all models in a congruence class, *H* is also constant across the congruence class.

The probability of obtaining a tree of size *n* ≥ 1 conditional upon the age of the stem lineage (*τ*_*o*_) and its survival to the present, denoted *P*_stem_(*n*), is given by the ratio *P* (*n*)/(1 − *E*(*τ*_*o*_)), i.e.:

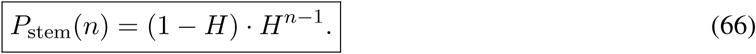

Since *H* is constant across a congruence class, the same also holds for *P*_stem_(*n*) for any *n*. The probability of obtaining a tree of size *n* ≥ 1 conditional upon the splitting of the root at age *τ*_*o*_ and the survival of its two daughter lineages, denoted *P*_root_(*n*), can be derived in a similar way, as follows. The probability that the two daughter lineages survive, conditional upon the split at age *τ*_*o*_, is given by the product:

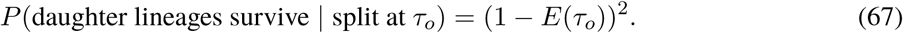

The probability that the two daughter lineages survive and the timetree has size *n* ≥ 1, conditional upon the split at age *τ*_*o*_, is given by the following sum of probabilities:

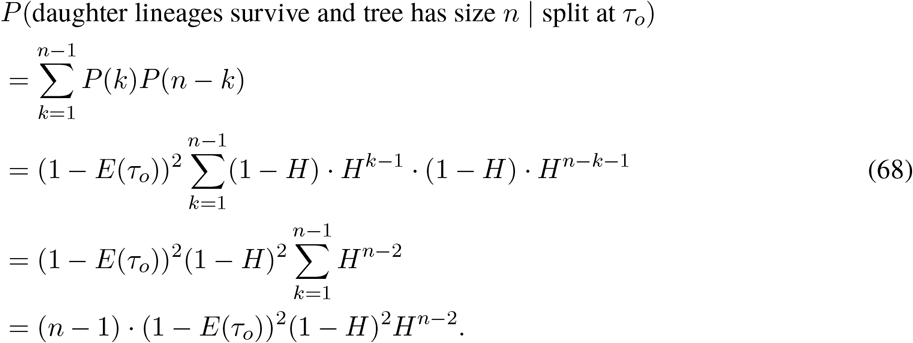

Dividing Eq. (68) by Eq. (67) yields the desired probability:

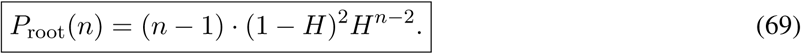

Since *H* is constant across the congruence class, the same also holds for *P*_root_(*n*).

##### S.1.8 The geometric nature of congruence classes

In the following, we provide a geometric interpretation of model congruence classes, by pointing out an analogy to the concept of object congruency in geometry. A basic background in abstract algebra is assumed.

In geometry, two objects are called congruent if they exhibit similar geometric properties, such as identical angles between corresponding lines and identical distances between corresponding points. More precisely, two geometric objects (sets of points in Euclidean space ℝ^*n*^) are called congruent if one set can be transformed into the other set by means of an isometry, i.e. a mapping that preserves distances between pairs of points (via translations, rotations, and/or reflections). Object congruency is a type of equivalence relation, and hence the set of models congruent to some focal object is an equivalence class. The set of all isometries is itself a group (known as “Euclidean group”) that acts on the set of geometric objects, and congruence classes of objects correspond to “orbits” under the action of isometries (8).

By analogy, two birth-death models are called “congruent” if they exhibit similar statistical properties in terms of their generated extant timetrees and LTTs (see main text and Supplement S.1). In fact, congruence classes can be interpreted as the orbits of a group of mappings acting on model space that preserve dLTTs (just as isometries preserve distances in Euclidean space). For technical reasons, we shall henceforth only consider the space of birth-death models (denoted 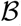) with strictly positive *λ*, *µ* and *ρ* and continuously differentiable *λ* and *µ* defined over some age interval [0, *τ*_*o*_] ⊆ ℝ. Let 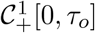 denote the set of all continuously differentiable real-valued strictly positive functions defined on the interval [0, *τ*_*o*_]. For any *S*_*o*_ ∈ (0, ∞) and any 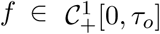, define 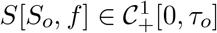 as the solution to the following initial value problem:

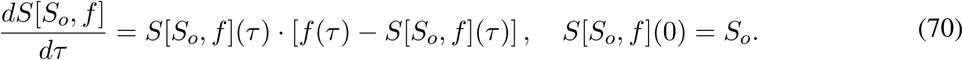

It is straightforward to verify that the solution to the above problem is given by:

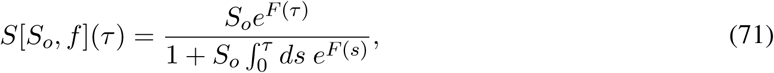

where we denoted:

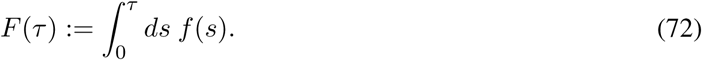

For any arbitrary *α* ∈ (0, ∞) and 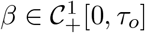, let 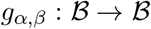 be a transformation of birth-death models defined as follows:

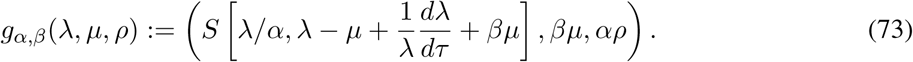

Note that *g*_*α,β*_ is dLTT-preserving, that is, it maps models to models within the same congruence class. Indeed, the variable

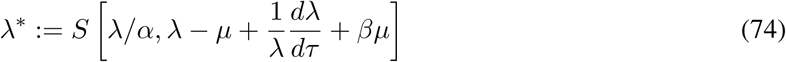

is exactly the speciation rate of a model with extinction rate 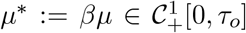 and sampling fraction *ρ*\*:= *αρ* ∈ (0, ∞), congruent to the original model (*λ, µ, ρ*). The set of all such transformations,

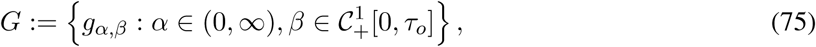

constitutes a group with group operation:

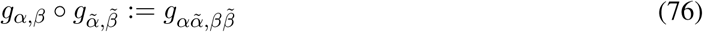

and identity element *g*_1,1_. The group *G* acts on the set of birth-death models, while preserving dLTTs. Abstractly, each mapping *g* ∈ *G* corresponds to an “isometric” transformation in model space that preserves the statistics of generated extant timetrees and dLTTs, in analogy to how rotations, translations or reflections preserve distances in Euclidean space.

Note that not all dLTT-preserving mappings defined on 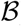 are members of *G*. It turns out, however, that *G* is large enough to completely generate congruence classes in 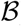. In other words, for any model 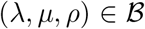, the orbit:

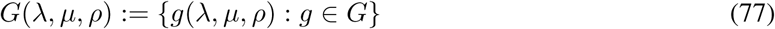

is exactly the congruence class of the model; indeed, for any congruent model 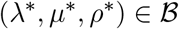 one can find a transformation *g*_*α,β*_ ∈ *G* such that (*λ*\*, *µ*\*, *ρ*\*) = *g*_*α,β*_(*λ, µ, ρ*), by choosing *α*:= *ρ*\*/*ρ* and *β*:= *µ*\*/*µ*.

### S.2 Why previous studies failed to detect model congruencies

In practice, reconstructions of *λ* and *µ* over time are typically performed by selecting among a limited set of allowed models, i.e., considering specific functional forms described by a finite number of parameters (9, 10, 11, 12, 13, 5). In these situations it is generally unlikely that the allowed model set intersects a given congruence class more than once (see Supplement S.3 for mathematical justification). For example, when considering only constant-rate birth-death models and assuming that *ρ* is fixed (as is usually the case; 14), each congruence class reduces to a single combination of *λ* and *µ*. Likelihood functions defined over a limited allowed model set thus generally don’t exhibit ridges associated with congruence classes, and may even exhibit a unique global maximum in the space of considered parameters, leaving the impression that *λ* and *µ* have been estimated close to their true values. Our findings suggest that this impression is almost certainly false. Instead, obtained estimates for *λ* and *µ* are almost always going to be a random outcome that depends on the particular choice of allowed models, such as the functional forms considered for *λ* and *µ*, and will be as close as possible to the congruence class of the truth rather than close to the truth itself. Unless one has reasons to prefer specific functional forms for *λ* and *µ* (e.g., based on a mechanistic macroevolutionary model; 15), fitted *λ* and *µ* are unlikely to resemble the true rates even if in principle the functional forms considered are flexible enough to resemble the true *λ* and *µ* (see Supplement S.6 for examples using simulations and real data).

By analogy, studies that test whether diversification dynamics are influenced by some environmental or geological variable *X* (e.g., temperature), either by testing for correlations between *X* and the estimated *λ* or *µ* (16, 17) or by fitting models in which *λ* or *µ* are explicit functions of *X* (18, 19, 20), will generally lead to unreliable conclusions. Indeed, specifying *λ* or *µ* as functions of *X* (e.g., assuming *µ* = *αX* + *β* and fitting the coefficients *α* and *β*) is essentially equivalent to choosing particular functional forms for *λ* or *µ*. Incidentally, evaluations of estimation methods based on simulations of the same limited model set as considered by the very estimation method evaluated (9, 11, 12, 13, 5), for example simulating trees with linearly varying *λ* and *µ* and then fitting models with linearly varying *λ* and *µ* to check if linear coefficients are accurately estimated, can generate the false impression that *λ* and *µ* can in general be reliably identified. Indeed, any given simulated model is typically going to be the sole representative of its congruence class that’s also in the method’s allowed model set.

### S.3 Typical model sets do not exhibit congruence ridges

In the following we explain why it is unlikely in practice that a limited set of allowed models (e.g., considered for maximum-likelihood estimation) will intersect any given congruence class more than once, and that it is especially unlikely that multiple intersections of a congruence class form a sub-manifold in parameter space (i.e., a “congruence ridge”). Consider a set of allowed models, parameterized through *n* independent parameters *q*_1_, .., *q_n_ ∈* ℝ, i.e. such that the speciation and extinction rates of a model are given as functions of age (*τ*) and the chosen parameters (**q** ∈ ℝ^*n*^):

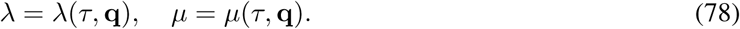

For simplicity, assume that the sampling fraction *ρ* is given (identifiability issues associated with uncertainties in *ρ* are already well known; 21, 22, 23, 24).

Now consider some particular choice of parameters, **q**, with corresponding PDR:

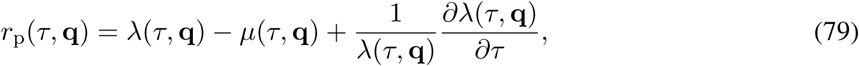

and present-day speciation rate *λ*(0, **q**). For any other choice of parameters **h** ∈ ℝ^*n*^, the corresponding model would be in the same congruence class as the first model if and only if *λ*(0, **h**) = *λ*(0, **q**) and *r*_p_(*τ*, **h**) = *r*_p_(*τ*, **q**) for all ages *τ ≥* 0, in other words *λ*(*⋅*, **h**) must be a solution to the initial value problem:

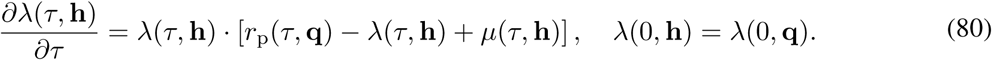

Unless the functional forms of *λ* and *µ* have been specifically designed for this purpose, it is generally unlikely that Eq. (80) will be satisfied for some **h** *≠* **q**.

A stronger argument for the low probability of congruence ridges can be made as follows. Suppose that **q** was part of a congruence ridge, i.e. a sub-manifold in parameter space belonging to the same congruence class. Then there must exist a curve in parameter space, i.e. a one-parameter function **h**: [−*ε, ε*] → ℝ^*n*^, passing through **q** (e.g., say **h**(0) = **q**), such that:

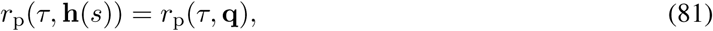

and such that:

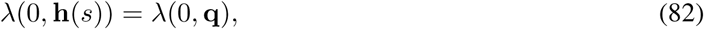

for all *s* ∈ [*−ε, ε*] and all *τ ≥* 0. Taking the derivative of Eq. (81) with respect to *s* at 0 yields:

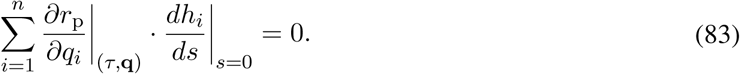

Denote 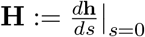 and 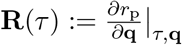. Then the condition in Eq. (83) can be written in vector notation:

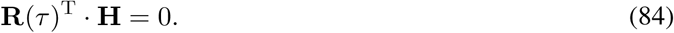

Note that **H** can be interpreted as the “velocity vector” along the ridge curve **h** at the point **q**, and hence condition (84) means that the ridge must move perpendicular to the direction of steepest descent of *r*_p_. Observe that condition (84) must be satisfied for all ages *τ* ≥ 0. Hence, for any arbitrary choice *τ*_1_*, τ*_2_*, .., τ_m_* ≥ 0, we obtain the following *m* linear equations that must be satisfied by **H**:

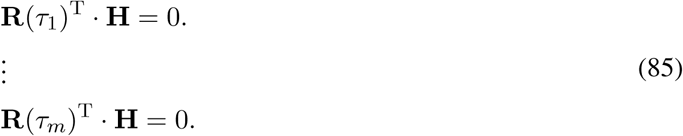

Unless the functional forms of *λ* and *µ* are specifically designed for this purpose, the system in Eq. (85) will almost certainly be over-determined if *m* is chosen sufficiently high (*m* ≫ *n*). Hence, in practice, for a chosen set of allowed models and a given point **q** in parameter space, a congruence ridge will almost never exist at that point.

## S.4 Interpreting the PDR

Given that the PDR is a composite quantity that depends on both *λ* and *µ* (Eq. 1), properly interpreting the estimated PDR in terms of actual speciation/extinction rates remains the responsibility of the investigator. Previous work has shown that the PDR can indeed yield valuable insight into diversification dynamics and can be useful for testing alternative hypotheses (6). For example, sudden rate transitions (e.g., due to mass extinction events) almost always lead to fluctuations in the PDR; thus, a relatively constant PDR over time would be indicative of constant — or only slowly changing — speciation and extinction rates.

The PDR can be used to obtain other useful variables. For example, it is straightforward to confirm that the PDR and the total diversity *N* satisfy the following relationship:

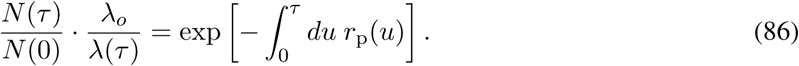

Observe that the left hand side of this equation, henceforth called deterministic “pulled normalized diversity” (dPND), corresponds to the ratio of deterministic total diversity at some age *τ* over the assumed present-day total diversity *N* (0), modulated by the factor *λ*_*o*_/*λ*(*τ*). Like the PDR, the dPND is the same for all models in a congruence class, and can be readily estimated from extant timetrees (Fig. S4C). As becomes apparent from Eq. (86), while the dPND can yield information on variations of past diversity, the amount of information depends on how well *λ* can be constrained a priori.

Another useful derived variable is the “pulled extinction rate”, or PER (6), defined as:

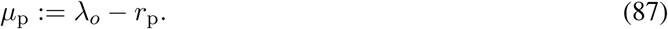

The PER is equal to the extinction rate *µ* if *λ* is time-independent, but differs from *µ* in most other cases. Note that calculating the PER requires knowing the present-day speciation rate *λ*_*o*_, which can be estimated from the timetree if the sampling fraction *ρ* is known (simply divide the estimated *ρλ*_*o*_ by *ρ*). The present-day PER is related to the present-day extinction rate as follows:

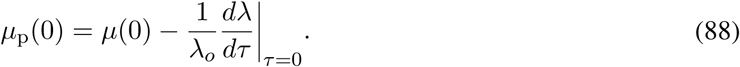

Observe that if the present-day speciation rate changes only slowly, the present-day PER will resemble the present-day extinction rate. Further, since *µ*(0) is non-negative, we can obtain the following lower bound for the exponential rate at which *λ* changes:

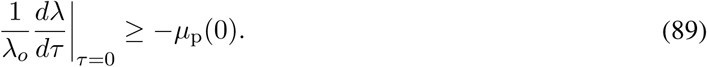

In particular, if the estimated *µ*_p_(0) is negative, this is evidence that *λ* is currently decreasing over time.

## S.5 Fitting congruence classes instead of models

The discussion in the main article revealed that speciation and extinction rates constitute partly interchangeable (and thus partly redundant) parameters that cannot be completely resolved from extant timetrees alone, no matter how large the dataset. Extant timetrees do, however, contain the proper information to estimate the pulled diversification rate *r*_p_ and *η*_*o*_ (recall that *η*_*o*_ = *ρλ*_*o*_), and may thus be used to at least identify the congruence class from which a tree was likely generated. In fact, for sufficiently large trees *r*_p_ and *η*_*o*_ can be directly calculated from the slope and curvature of the tree’s LTT (6). Since each congruence class corresponds to a unique *r*_p_ and *η*_*o*_, the *r*_p_ and *η*_*o*_ can be used to parameterize the space of congruence classes; on this space the likelihood function no longer exhibits the highly problematic ridges seen in the original model space. We thus suggest describing birth-death models in terms of *r*_p_ and *η*_*o*_, rather than *λ* and *µ*, when fitting models to timetrees. Since the likelihood function can be expressed directly in terms of *r*_p_ and *η_o_* (Supplement S.1.6), such a parameterization is suitable for maximum-likelihood or Bayesian estimation methods. Reciprocally, since every given *r*_p_ and *η*_*o*_ correspond to a unique and non-empty congruence class (as shown in the main article), any *r*_p_ and *η*_*o*_ estimated from an extant timetree will represent at least one biologically meaningful scenario. It is thus possible to directly fit congruence classes, rather than individual models, via maximum-likelihood. A similar reasoning can also be applied to the pulled speciation rate *λ*_p_, which provides an alternative representation of congruence classes.

To demonstrate this approach, we created software for fitting *r*_p_ and *η*_*o*_ to extant timetrees via maximum likelihood. The code is integrated into the R package castor (25) as function `fit_hbd_pdr_on_grid`. The function accepts as input an extant timetree, and an arbitrary number of discrete ages at which to estimate *r*_p_, assuming *r*_p_ varies linearly or polynomially between those ages. The function also accepts optional lower and upper bounds for the fitted *r*_p_ and/or *η*_*o*_. The code then maximizes the likelihood of the tree, given by Eq. (56) in the Supplement, by iteratively refining the *r*_p_ values on the age grid and/or *η*_*o*_. Optionally, one can limit the evaluation of the likelihood function to a smaller “truncated” age interval than covered by the tree, i.e. some age interval [0, *τ*^*\**^], where *τ*^*^ may be smaller than the root age. This may be useful for avoid estimation errors towards older ages due to a small number of lineages in the tree. The likelihood formula for the “truncated” case can be easily obtained by assuming that the tree is split into multiple sub-trees, each originating at the truncation age, and considering each sub-tree an independent realization of the same birth-death process and subject to the same sampling fraction *ρ*. To avoid non-global local optima, the fitting can be repeated multiple times, each time starting at random start values for the fitted parameters, and the best fit among all repeats is kept. We also developed similar computer code for fitting the pulled speciation rate *λ*_p_ to extant timetrees, implemented as function `fit_hbd_psr_on_grid` in the R package castor.

Supplemental Fig. S4 shows an example where either the *r*_p_ or *λ*_p_ were accurately fitted to an extant timetree, simulated under a birth-death scenario subject to an early rapid radiation event and followed by a mass extinction event. In this example, we limited fitting to ages where the LTT was over 500 lineages (i.e., *M* (*τ ^*^*) = 500), and repeated the fitting 100 times to avoid non-global local optima.

## S.6 Fitting birth-death models to trees yields unreliable results

To illustrate the identifiability issues discussed in the main article, we simulated and analyzed two massive extant timetrees (~ 114,000 and ~785,000 tips) via a birth-death process, subject to a mass extinction event (both trees) and a rapid radiation event (second tree). Instead of fitting models of the exact same functional form as used in the simulations, we fitted generic piecewise-linear curves for *λ* and *µ* that could in principle take various alternative shapes (including approximately the shapes used for the simulations), and visually compared the estimated profiles to their true profiles (Supplemental Figs. S5A–F). Specifically, we fitted *λ* and *µ* at multiple discrete time points, treating the rates at each time point as free parameters, while assuming a known *ρ*. Despite the enormous tree sizes, and despite the fact that the fitted models reproduced the trees’ LTTs and the true *r*_p_ extremely well (Supplemental Figs. S5A,C,D,F), the estimated *λ* and *µ* were far from their true values and even exhibited spurious trends (Supplemental Figs. S5BE). This is consistent with our expectation that the particular combination of fitted *λ* and *µ* is essentially a random pick from the periphery of the true process’s congruence class. In contrast, when we fixed *µ* to its true profile, *λ* was accurately estimated (Supplemental Fig. S2), consistent with the expectation that any given *µ* and *ρ* fully determine the corresponding *λ* in the congruence class.

We also examined a large extant timetree of 79,874 seed plant species (Supplemental Fig. S3) published by Smith *et al.* (26), and estimated *λ* and *µ* over the last 100 million years using two alternative approaches (methods details in Supplement S.7). In the first approach, we fitted generic piecewise-linear curves for *λ* and *µ*, similarly to the previous example. In the second approach, we fitted parameterized time curves for *λ* and *µ* that included an exponential as well as a polynomial term (5). Even though both approaches yielded similar estimates for *r*_p_, and both accurately reproduced the tree’s LTT, they yielded markedly different *λ* and *µ* (Supplemental Figs. S5D–F). This observation is consistent with our argument that, depending on the precise set of models considered, the estimated *λ* and *µ* will generally be a random pick from the underlying (true or close-to-true) congruence class.

To illustrate our point that common model selection approaches such as minimizing the Akaike Information Criterion (AIC) (27) cannot resolve the identifiability issues discussed, we also fitted a series of models of variable complexity to a massive timetree of 1,000,000 tips. The tree was simulated based on origination and extinction rates of marine invertebrate genera, previously estimated from marine invertebrate fossil data (28) (Fig. 2D in the main article). We fitted two types of models: piecewise constant models and piecewise linear models. In piecewise constant models (sometimes also referred to as “birth-death-shift” models; 13) the rates *λ* and *µ* have constant values within discrete time intervals, with every time interval exhibiting distinct *λ* and *µ*. In piecewise linear models *λ* and *µ* vary linearly between discrete time points. For both model types we considered various temporal grid sizes, ranging from 5 up to 15 grid points, thus including sufficient model complexity for approximating the true rates. In all cases the time grid points where located at equidistant intervals between the present and the tree’s root. For each model type (piecewise constant or piecewise linear) and for each grid size we estimated the free parameters (either the rates within each interval, or the rates at each grid point, respectively) via maximum likelihood using the function `fit_hbd_model_on_grid` in the R package `castor`. Only the most recent 100 million years were considered for fitting, in order to focus estimations on times with greater lineage density in the simulated tree (towards the root estimated rates will be inaccurate regardless of the arguments presented in this paper). Fitting was repeated 20 times with random start parameters to avoid local non-global optima. Among each model type, we then kept the maximum likelihood model with smallest AIC value, shown in Supplemental Fig. S6. As expected, estimated rates were highly inaccurate and missed important features, despite the fact that we were using a massive tree of 1,000,000 tips, and the fact that the tree’s LTT was almost perfectly matched by the models’ dLTTs.

## S.7 Fitting birth-death models to seed plants

An extant timetree of 79,874 seed plant species, constructed using GenBank sequence data with a backbone provided by Magallón *et al.* (29), was obtained from the Supplemental Material published by Smith *et al.* (26, tree “CBMB”). The tree is shown in Supplemental Fig. S3. The sampling fraction was calculated based on the tree size and the number of extant seed plant species estimated at 422,127 by Govaerts (30). As mentioned in Supplement S.6, two approaches were used to fit *λ* and *µ* over time. In the first approach, *λ* and *µ* were allowed to vary independently at 8 discrete and equidistant time points (assuming piecewise linearity between grid points) and were estimated via maximum-likelihood using the function `fit_hb_model_on_grid` in the R package castor (25) (options `“condition=‘stem’, relative_dt=0.001”`). Fitting was repeated 100 times using random start parameters to avoid local non-global optima in the likelihood function. The PDR was then estimated from the fitted *λ* and *µ* using the formula in Eq. (1) and using central finite differences to calculate derivatives on the time grid. In the second approach, *λ* and *µ* were assumed to be of the following general functional forms:

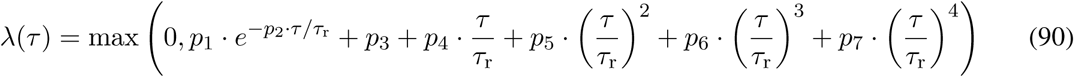

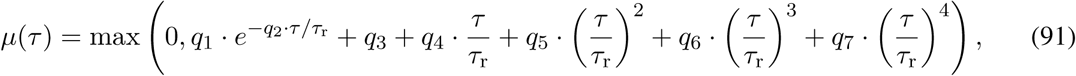

where *τ*_r_ is the age of the root and *p*_1_, .., *p*_7_, *q*_1_, .., *q*_7_ are fitted parameters. Parameters were fitted using the `castor` function `fit_hbd_model_parametric` (options `“condition=‘stem’, relative_dt=0.001, param_min=-100, param_max=100”`). As in the first approach, fitting was repeated 100 times to avoid local non-global optima. In both approaches, the likelihood only incorporated branching events at ages between 0 and 130 Myr, since the LTT and any parameter estimates become much less reliable at older ages.

## S.8 Supplemental figures

**Figure S1:**
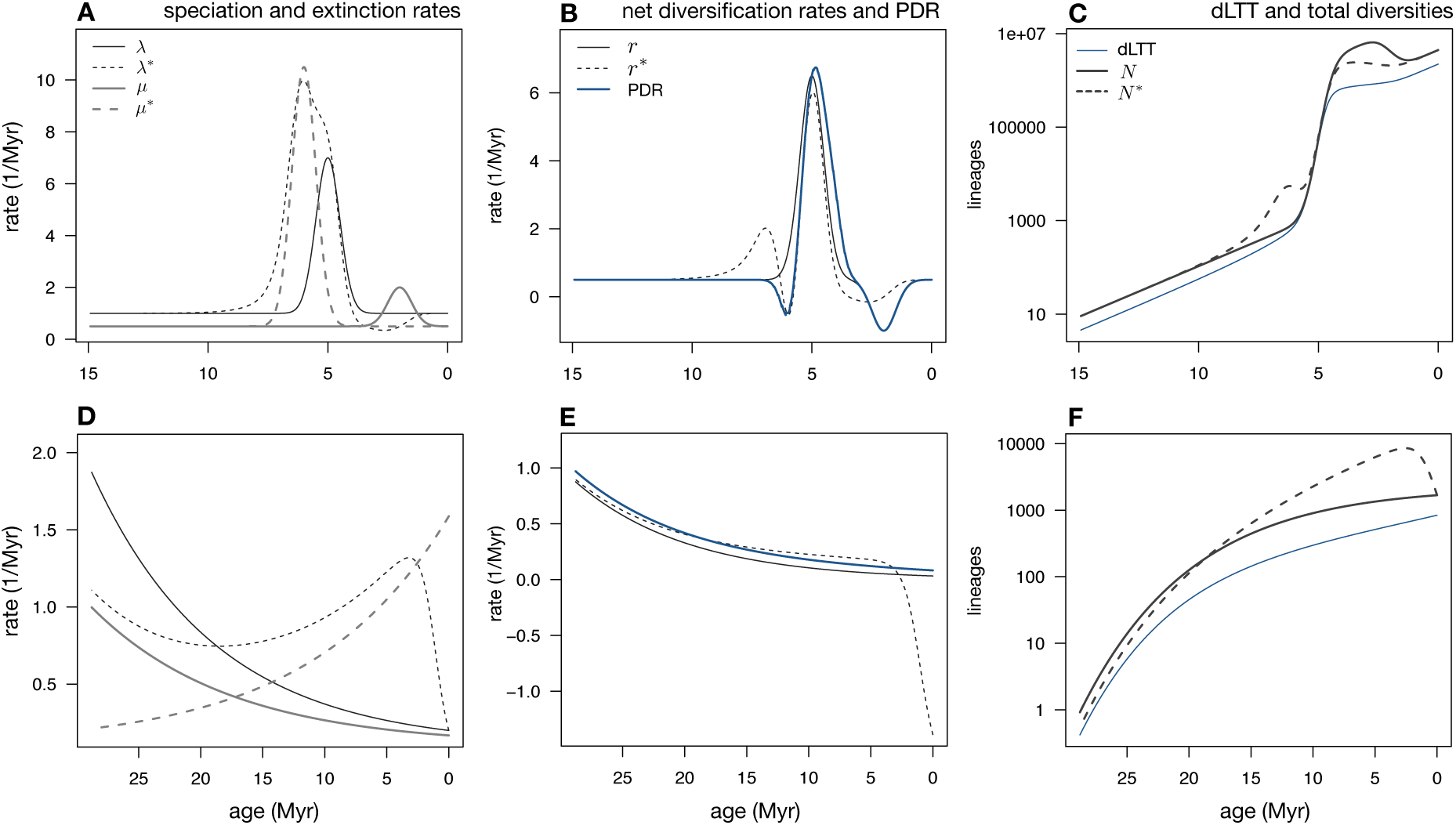
Examples of congruent birth-death processes. (A–C) Example of two congruent yet markedly different birth-death models. Both models exhibit a temporary spike in the extinction rate and a temporary spike in the speciation rate, however the timings of these events differ substantially between the two models. Both models exhibit the same dLTT and the same pulled diversification rate *r*_p_, and would yield identical likelihoods for any given extant timetree. (A) Speciation rates (*λ* and *λ^*^*) and extinction rates (*µ* and *µ^*^*) of the two models, plotted over time. Continuous curves correspond to the first model, dashed curves to the second model. (B) Net diversification rates (*r* and *r^*^*) and pulled diversification rate *r*_p_ of the two models. (C) Deterministic LTT (dLTT) and deterministic total diversities (*N* and *N ^*^*) predicted by the two models. (D–F) Another example of two congruent models. In the first model, the speciation and extinction rates both decrease exponentially over time, whereas in the second model the extinction rate increases exponentially over time and the speciation rate exhibits variable directions of change over time. In all models the sampling fraction is *ρ* = 0.5.

**Figure S2:**
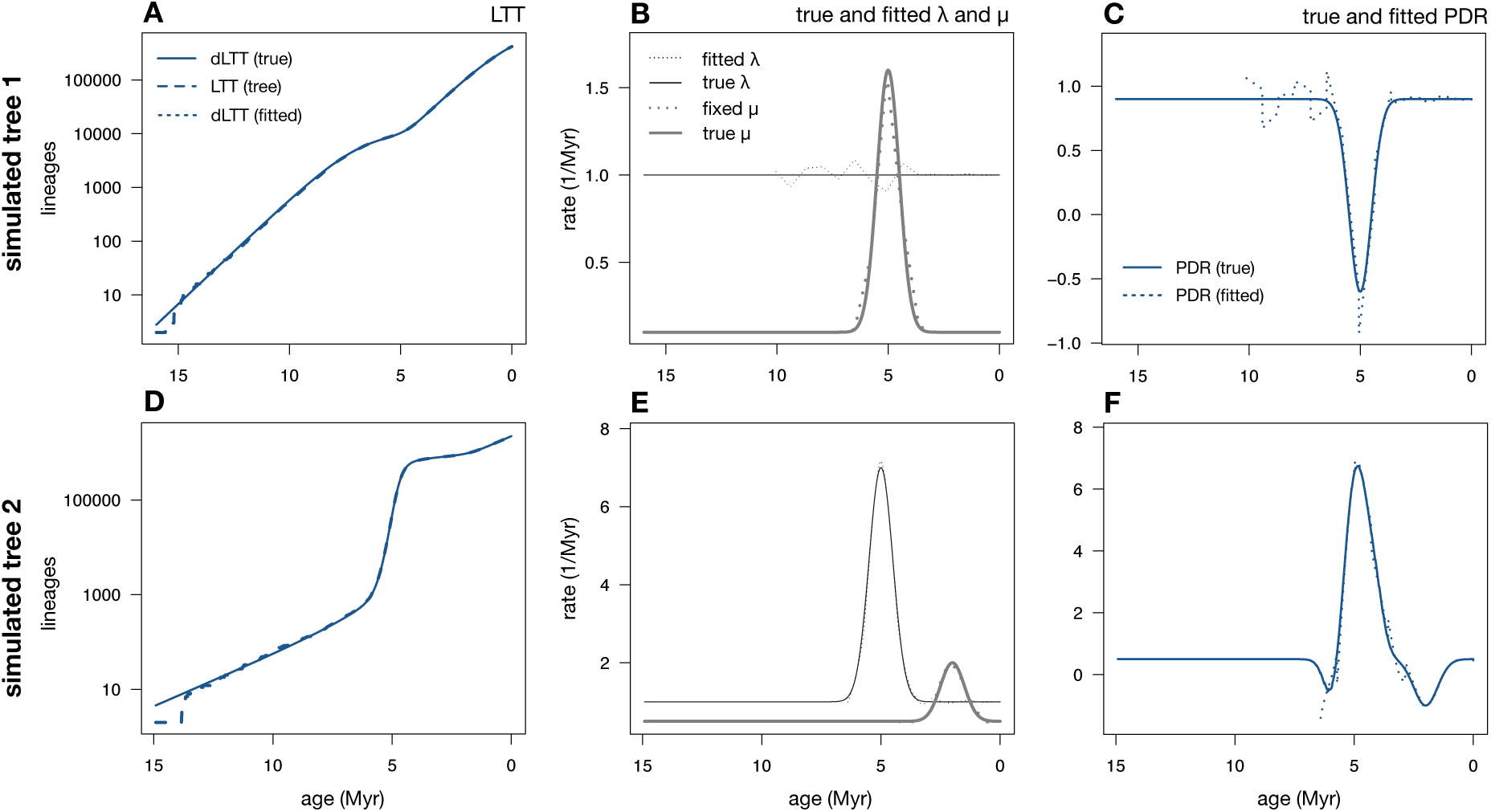
Estimating *λ* when *µ* and *ρ* are fixed. (A–C) Example analysis of a simulated extant timetree (~114,000 tips), exhibiting a mass extinction event at ~5 Myr before present. A birth-death model was fitted while fixing *µ* and *ρ* to their true values; *λ* was fitted at 15 discrete time points. (A) Lineages through time curve (LTT) of the generated tree (long-dashed curve), deterministic LTT (dLTT) of the true model that generated the tree (continuous curve), and dLTT of a maximum-likelihood fitted model (short-dashed curve). The fitted dLTT is practically identical to the true dLTT and is thus covered by the latter. (B) True speciation and extinction rates (continuous curves), along with the fitted speciation rate and fixed extinction rate (dashed curves). (C) Pulled diversification rate (PDR) of the true model (*r*_p_, continuous curve), compared to the PDR of the fitted model (dashed curve). (D–F) Example analysis of a simulated extant timetree (~785,000 tips), exhibiting a rapid radiation event at ~5 Myr before present and a mass extinction event at ~2 Myr before present. A birth-death model was fitted similarly to the previous example, and D–F are analogous to A–C. In both cases, rate estimation was restricted to ages where the LTT included at least 500 lineages.

**Figure S3:**
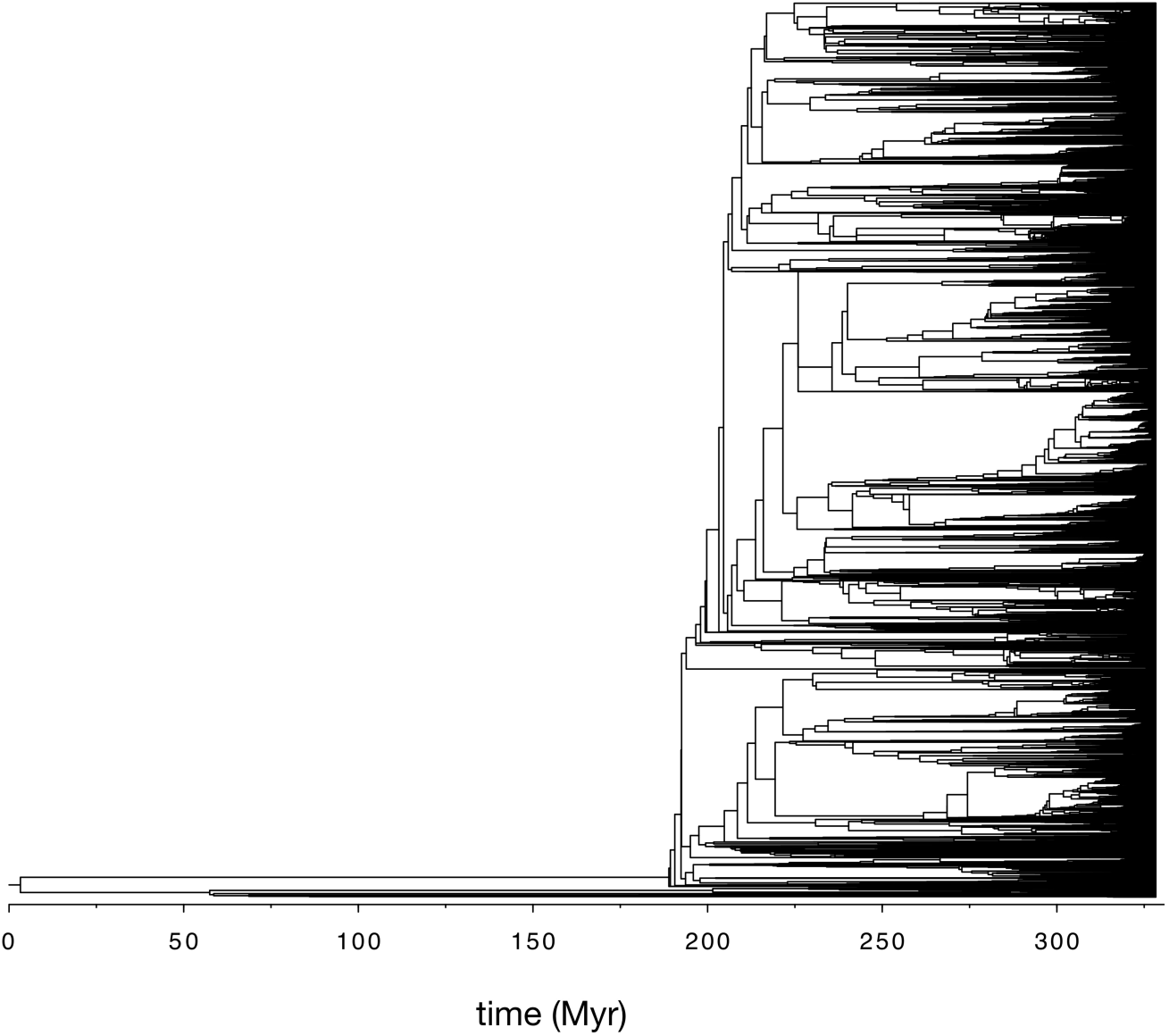
Seed plant tree. Extant timetree of 79,874 seed plant species, discussed in the main article. The tree was constructed and made available by Smith *et al.* (26) (tree “GBMB”).

**Figure S4:**
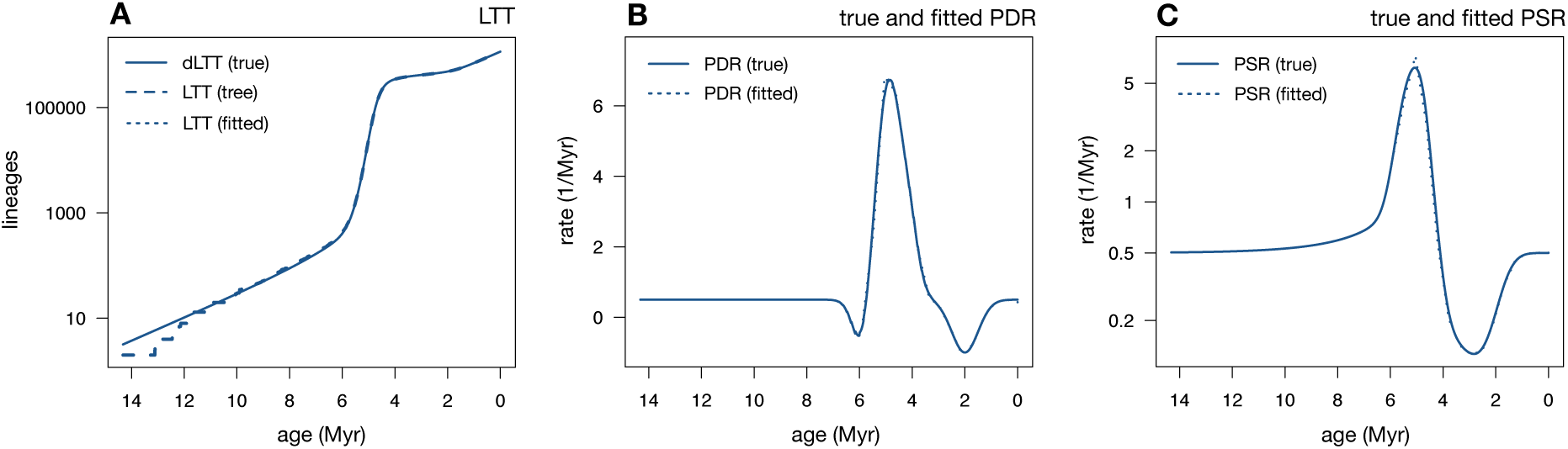
Fitting congruence classes instead of models. Analysis of an extant timetree generated by a birth-death model, exhibiting a temporary rapid radiation event about 5 Myr before present and a mass extinction event about 2 Myr before present. A congruence class was fitted to the timetree either in terms of the pulled diversification rate (PDR, *r*_p_) and the product *ρλ_o_*, or in terms of the pulled speciation rate (PSR, *λ*_p_). (A) Lineages through time curve (LTT) of the tree (long-dashed curve), together with the deterministic LTT (dLTT) of the true model (continuous curve) and the dLTT of the fitted congruence classes (short-dashed curve); in both cases the fitted dLTT was virtually identical to the true dLTT, and is thus completely covered by the latter. (B) PDR of the true model (continuous curve), compared to the fitted PDR (short-dashed curve). (C) PSR of the true model (continuous curve), compared to the fitted PSR (short-dashed curve). The PDR and PSR were fitted via maximum-likelihood using the R package `castor` (25), allowing the PDR or PSR to vary freely at 15 discrete equidistant time points (Supplement S.5).

**Figure S5:**
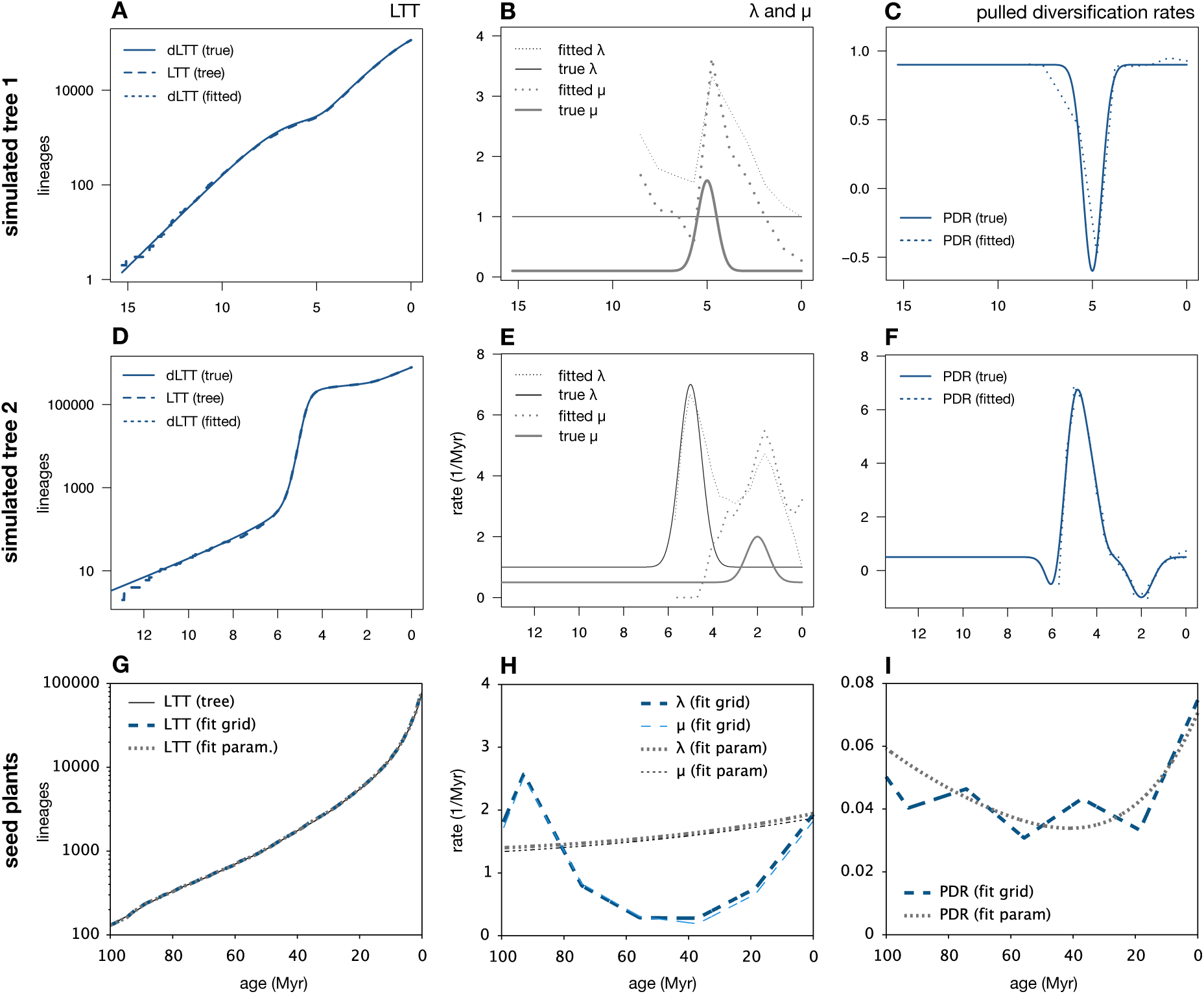
Identifiability issues persist in large trees. (A–C) Diversification analysis of a timetree (~ 114,000 tips) simulated from a birth-death process exhibiting a mass extinction event at ~5 Myr before present. (A) Lineages through time curve (LTT) of the generated tree (long-dashed curve), deterministic LTT (dLTT) of the true model that generated the tree (continuous curve), and dLTT of a maximum-likelihood fitted model (short-dashed curve). The fitted dLTT is practically identical to the true dLTT and is thus covered by the latter. (B) True speciation and extinction rates (continuous curves), compared to fitted speciation and extinction rates (dashed curves). Observe the great disagreement between the fitted and true *λ* and *µ*, despite the fact that the allowed model set could in principle approximate the true rates reasonably well. (C) Pulled diversification rate (PDR) of the true model (continuous curve), compared to the PDR of the fitted model (dashed curve). (D–F) Diversification analysis of a timetree (~785,000 tips) simulated from a birth-death process exhibiting a rapid radiation event at ~5 Myr before present and a mass extinction event at ~2 Myr before present. Sub-figures D–F are analogous to A–C. Again, observe the great disagreement between the fitted and true *λ* and *µ*, despite the fact that the allowed model set could in principle approximate the true rates reasonably well. See Supplemental Fig. S2 for fitting results when *µ* is fixed to its true value. (G–I) Diversification analyses of an extant timetree of 79,874 seed plant species (26), performed either by fitting *λ* and *µ* on a grid of discrete time points or by fitting the parameters of generic polynomial/exponential functions for *λ* and *µ*. (G) LTT of the tree, dLTT of the grid-fitted model and dLTT of the fitted parametric model. (H) Speciation and extinction rates predicted by the grid-fitted model or the fitted parametric model. (I) PDR predicted by the grid-fitted model and the fitted parametric model.

**Figure S6:**
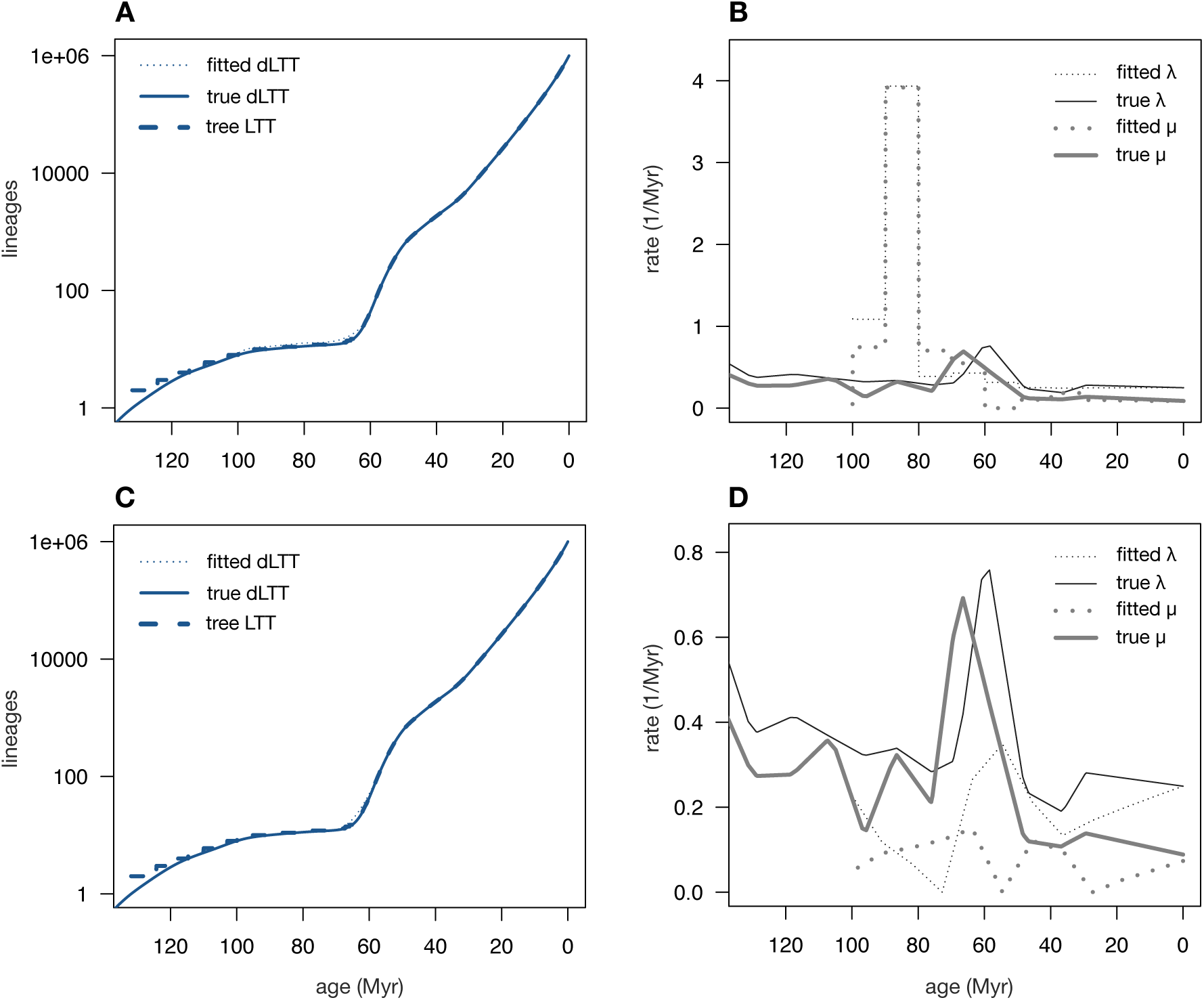
Identifiability issues cannot be resolved with AIC. Maximum-likelihood birth-death models fitted to a tree comprising 1,000,000 tips, simulated based on the origination and extinction rates of marine invertebrate genera estimated from fossil data (28). Top row: Maximum-likelihood-fitted piecewise constant model (also known as birth-death-shift model), with grid size (*N* = 11) chosen by minimizing the AIC. Bottom row: Maximum-likelihood-fitted piecewise linear model, with grid size (*N* = 12) chosen by minimizing the AIC. Left column: dLTTs of the fitted models compared to the true dLTT and the tree’s LTT. Right column: Fitted speciation and extinction rates, compared to the true rates used to generate the tree. Observe that in both cases the maximum-likelihood models poorly reflect the true rates despite a near-perfect match of the LTT, even when the complexity of the models was optimized based on the AIC. For Methods details see Supplement S.6.

